# A Well-Characterized Polycistronic-Like Gene Expression System in Yeast

**DOI:** 10.1101/2022.05.23.493076

**Authors:** Minakshi Mukherjee, Zhen Q. Wang

## Abstract

Efficient expression of multiple genes is critical to yeast metabolic engineering because of the increasing complexity of engineered pathways. A yeast polycistronic expression system is of particular interest because one promoter can drive the expression of multiple genes. Polycistronic expression thus requires less genetic material and minimizes undesirable recombination due to repeated use of the same promoter. It also decreases the number of DNA parts necessary for cloning a multi-gene construct. 2A viral peptides enable the co-translation of multiple proteins from one mRNA by ribosomal skipping. However, the wide adaptation of this strategy for polycistronic-like gene expression in yeast awaits in-depth characterizations. Additionally, a one-step assembly of such a polycistronic-like system is highly desirable. To this end, we have developed a MoClo compatible 2A peptide-based polycistronic-like system capable of expressing multiple genes from a single promoter in yeast. Characterizing the bi-, tri, and quad-cistronic expression of fluorescent proteins showed high cleavage efficiencies of three 2A peptides. The expression level of each protein decreases as it moves away from the promoter. Additionally, the impact of a C-terminal 2A peptide on protein function is dependent on the protein sequence. Applying the polycistronic-like system for geraniol production resulted in similar or higher titers compared to a conventional monocistronic construct. In summary, this highly-characterized polycistronic-like gene expression system is another tool to facilitate multi-gene expression in yeast.

## Introduction

Baker’s yeast (*Saccharomyces cerevisiae*) is widely used in metabolic engineering for renewable production of commodity and specialized chemicals^*1*^. The increasing lengths and complexity of engineered metabolic pathways in yeast require a highly efficient and controllable expression of multiple genes^*2-4*^. The recent development of new synthetic genetic parts has greatly enriched tools available for yeast gene expression^*5-14*^. However, gene expression in yeast is monocistronic such that each gene requires a separate promoter and terminator. As a result, multiple promoters and terminators are required to clone multi-gene constructs. This requirement often results in unusually large plasmids since the sizes of each promoter and terminator are considerable. Additionally, repetitive use of the same promoter increases unintended homologous recombination, while using different promoters causes uneven gene expression. Therefore, it will be of great advantage to develop a polycistronic-like system in yeast so that a single promoter will drive the expression of multiple genes.

2A viral peptides are viral oligopeptides of 20-30 amino acids, which “self-cleaves” a polypeptide into two separate proteins during translation^*15-17*^. Mechanistic studies showed that instead of a peptidyl cleavage, the 2A peptide mediates a highly unusual translation termination independent of the stop codon at the C-terminal conserved D(V/I) EXNPGP motif^*18, 19*^. Specifically, when the G of the D(V/I) EXNPG is at the P site in the ribosome, the 2A peptide: ribosome complex changes the fine structure to discriminate against the peptidyl transfer of the incoming prolyl-tRNA^Pro^. The ribosome thus stalls, releasing the first protein ending with the 2A peptide up to glycine. Then the ribosome resumes translation of the second peptide with proline as the first amino acid. Occasionally the ribosomal skipping does not happen, thus resulting in an “uncleaved” protein (Figure 1A). The highly efficient self-processing property and the small size render the 2A peptide an ideal tool to develop a polycistronic-like expression system in yeast.

**Figure 1.**
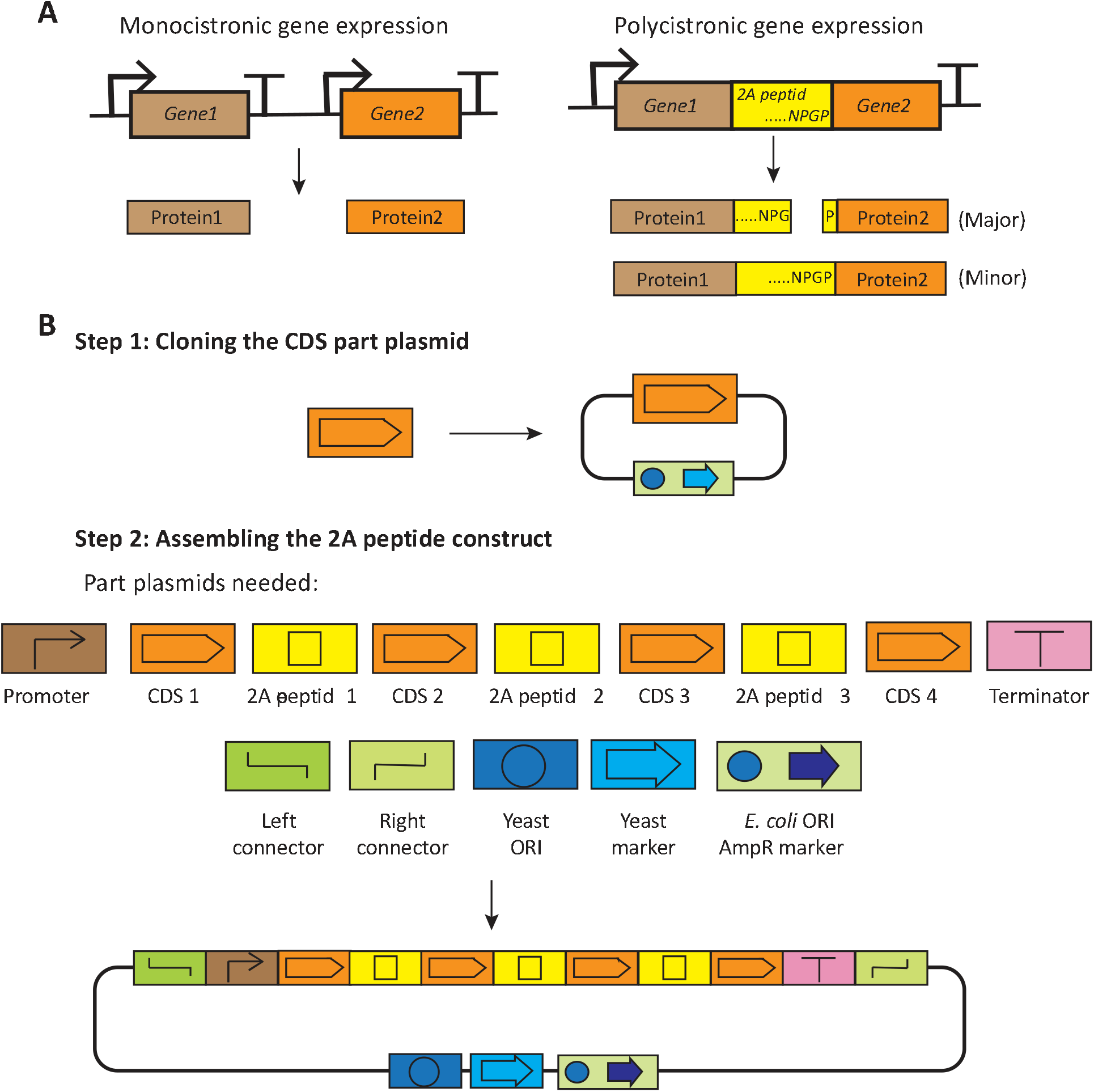
The 2A peptide-based polycistronic expression system in yeast. (A) Schematic of the canonical monocistronic and the 2A peptide-based polycistronic gene expression system. (B) Assembly of the 2A peptide-based polycistronic construct. Step 1: Cloning the gene of interest into the entry vector using the BsmbI (or Esp3I) enzyme to create the CDS part plasmid. Step 2: Assembly of part plasmids into a TU level vector using BsaI enzyme. The resulting polycistronic construct can be transformed into yeast. ORI: origin of replication; AmpR: ampicillin resistant gene.

2A peptides have been applied to express two or more genes in animals for antibody production and fluorescent cell imaging^*20-29*^. It has also been utilized to produce specialized metabolites in plants^*30-37*^ and fungus^*38-41*^. Recently, a pioneering work by Souza-Moreira et al. (2018)^*41*^ calculated cleavage efficiencies of twenty-two 2A peptides individually linking two fluorescent proteins in yeast. However, the 2A peptide system still requires further characterization before it can be widely adapted in yeast metabolic engineering. Questions remaining include: how reliable are 2A peptides when expressing more than two genes? How does the positioning of each gene in a 2A peptide construct affect its expression? Does the C-terminal 2A peptide affect the molecular function of the protein? Does the cleavage efficiency of a 2A peptide vary in different yeast strains? How does the 2A peptide construct compare to the traditional monocistronic expression system in terms of product yield? Is there a facile cloning method that allows the seamless assembly of multiple genes in a 2A peptide construct?

To address these questions, we first developed a Golden Gate compatible 2A polycistronic-like expression system that builds on the existing yeast MoClo kit developed by Lee et al. (2015)^*14*^. This system enables a one-step assembly of up to four genes separated by three 2A peptides. We used this system to characterize bi-, tri-, and quad-cistronic constructs using fluorescent proteins. Interestingly, we found that the 2A peptide tail negatively affects protein function but to a different extent in different proteins. The four 2A peptides characterized behaved similarly in four commonly used yeast strains. The position of each gene in a 2A peptide construct does influence its expression. Additionally, a side-by-side comparison of the conventional monocistronic expression system and the 2A peptide expression system was performed using a four-gene pathway for geraniol production. The assembled polycistronic constructs yielded a similar or higher amount of geraniol than their monocistronic counterparts.

## Results and Discussion

### Assembly of polycistronic-like expression constructs in yeast

We have developed a bottom-up approach to assemble 2A peptide-based polycistronic constructs building on the existing yeast MoClo toolkit^*14*^. In brief, the cloning process requires the assembly of part plasmids into a transcription unit (TU) level plasmid. The polycistronic constructs developed can assemble up to four coding sequences (CDS) linked by 2A peptides in a TU level construct, allowing the co-expression of four genes from a single promoter (Figure 1B). All of the part plasmids, except those containing CDSs, are available in the Addgene repository (kit #:1000000061). The codon-optimized 2A peptide part plasmids will be deposited to Addgene too. The users only need to clone their genes of interest into the entry vector followed by a one-step Golden Gate assembly into the TU level multi-gene construct. The fluorescent proteins in the entry vector greatly facilitate screening the correct constructs. It is routine to obtain the correctly assembled quad-cistronic construct in one attempt. The cloning protocol is detailed in Supplementary Figures 1-6 and Supplementary Tables 1-3.

This MolClo system greatly simplified the cloning of 2A peptide-linked genes. Firstly, the 2A peptide part plasmids have been sequence-verified; this eliminates the chance of mutations if they are synthetic oligonucleotides. Secondly, the Golden Gate protocol allows the simultaneous assembly of multiple genes in one step. The 2A peptide-based construct is at the TU level; there is no need to assemble a multi-gene plasmid, which is required for cloning conventional monocistronic constructs.

### Characterization of the bi-cistronic constructs

To characterize the expression of functional proteins in a bi-cistronic construct, we assembled four constructs with TagRFP-T as the first gene and mTurquoise2 as the second gene linked by a 2A peptide (Figure 2A). A GSG linker was placed between the first gene and the 2A peptide to maximize the cleavage efficiencies of the 2A peptides^*42*^ and minimize their potential impact on the function of the first protein. The four 2A peptides examined are E2A, O2A, P2A, and T2A. They are chosen based on a previous screening in yeast and the wide usage in other organisms^*20, 22, 24, 30, 31, 42*^. Fluorescence of both TagRFP-T and mTurquoise2 were detected, indicating the bi-cistronic system is functional (Figure 2B). Interestingly, TagRFP-T as the first gene was expressed at a lower level (43-80% of the control) than mTurquoise2, the second gene (72-78% of the control). One reason could be that the additional C-terminal 2A peptide of the first protein decreased fluorescence, despite a GSG linker. However, increasing linker lengths or switching to other linkers^*43*^ did not increase the fluorescence of the first protein (Supplementary Figure 7). Western blot using a C-terminal anti-his antibody showed high cleavage efficiency (∼95%) for E2A, O2A, and P2A. The cleavage efficiency of T2A was only 30% (Figure 2C).

**Figure 2.**
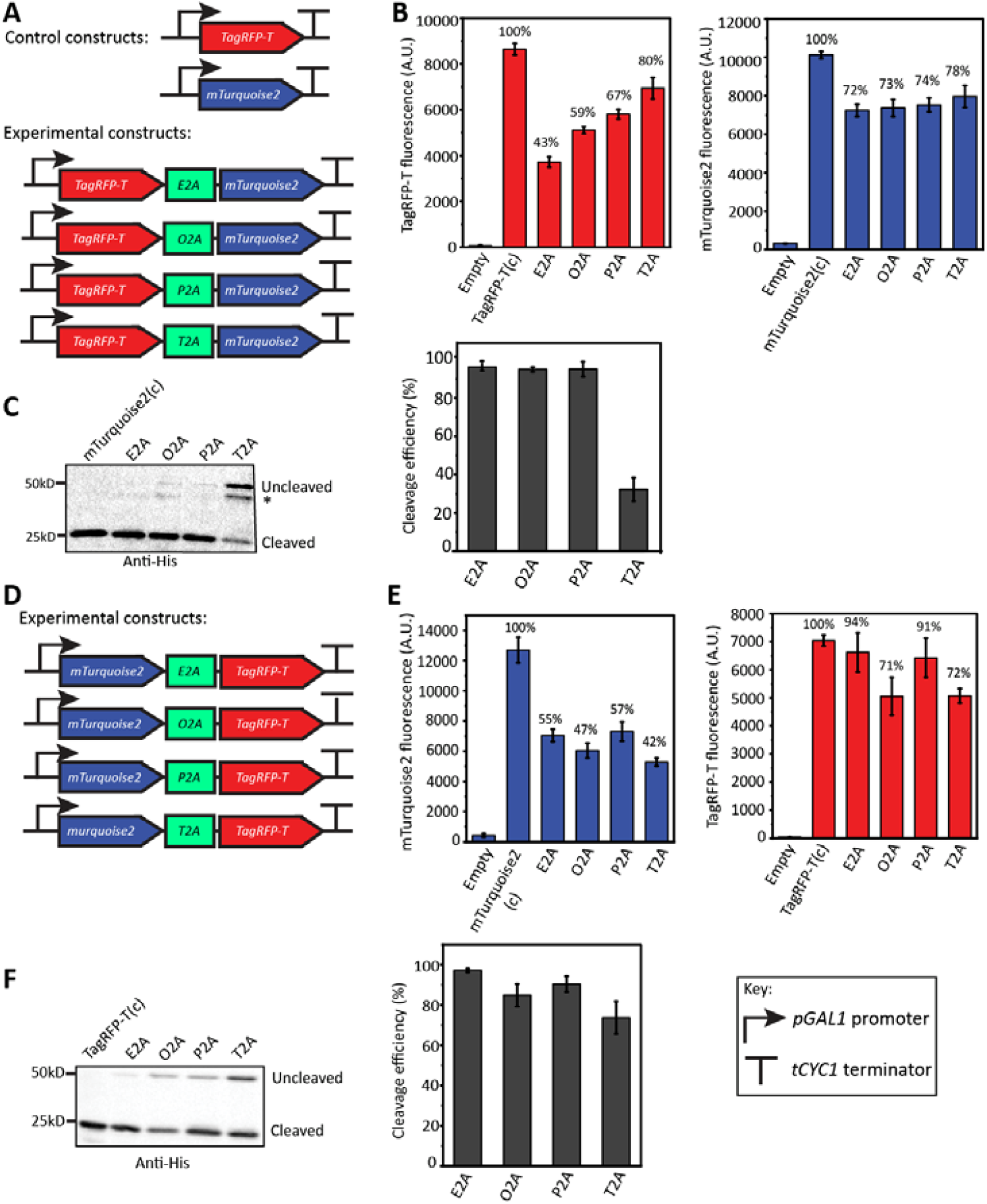
Characterization of bi-cistronic constructs. (A) Control monocistronic and experimental bi-cistronic constructs expressed under the *GAL1* promoter. E2A: 2A peptide from the Equine rhinitis B virus; O2A: 2A peptide from the Operophtera brumata cypovirus-18; P2A: 2A peptide from the Porcine teschovirus-1; T2A: 2A peptide from the Thosea asigna virus. (B) Normalized fluorescence over OD_600_ of TagRFP-T and mTurquoise2 in yeast strain BY4741. (C) Western blot to determine the cleavage efficiencies of the four 2A peptides. Anti-his antibody detects the 6*His tag at the C-terminus of mTurquoise2 in both control and experimental constructs. The asterisk indicates a major byproduct commonly observed in fluorescent proteins. Cleavage efficiencies (%) of the four 2A peptides are shown in the histogram. (D) The flipped bi-cistronic constructs with the positions of mTurquoise2 and TagRFP-T switched. (E) Normalized fluorescence of TagRFP-T and mTurquoise2 in the flipped constructs. (F) Western blot to determine the cleavage efficiency of the four 2A peptides in the flipped constructs. Data represent the average ± SD of three independent biological replicates.

To determine if the bi-cistronic expression is influenced by the two gene sequences flanking the 2A peptide, we flipped the construct so that mTurquoise2 was the first gene and TagRFP-T, the second (Figure 2D). Similar results were observed in the flipped constructs. mTurquoise2 as the first gene (42-57% of the control) was expressed significantly lower than the second gene (71-94% of control) (Figure 2E). The cleavage efficiencies for the four 2A peptides were similar to the original reporter constructs, with T2A being the least efficient. Thus, the expression and cleavage efficiency of bi-cistronic constructs does not seem to be affected by specific gene sequences.

### Effects of 2A peptides on protein function and strain dependence

Because the first fluorescent gene expression was consistently lower than its monocistronic control, we investigated if the 2A peptide interferes with protein function. To this end, we compared the function of fluorescent proteins and β-galactosidase lacZ with or without a C-terminal 2A peptide (Figure 3). Surprisingly, all three fluorescent proteins with a C-terminal E2A, including mTurquoise2, TagRFP-T, and mKate2, showed dramatically decreased fluorescence (2-9% of the control) (Figure 3B). However, lacZ with the C-terminal E2A only had a slight decrease of the β-galactosidase activity compared to the wildtype version (Figure 3C). Since fluorescent proteins and β-galactosidase are structurally disparate^*44, 45*^, the impact of the C-terminal 2A peptide on these two classes of proteins are drastically different. Therefore, how 2A peptide interferes with protein function varies from gene to gene. One may add a protease sequence between the first gene and the 2A peptide to mitigate the negative impact of the 2A peptide on the protein function^*28, 29*^.

**Figure 3.**
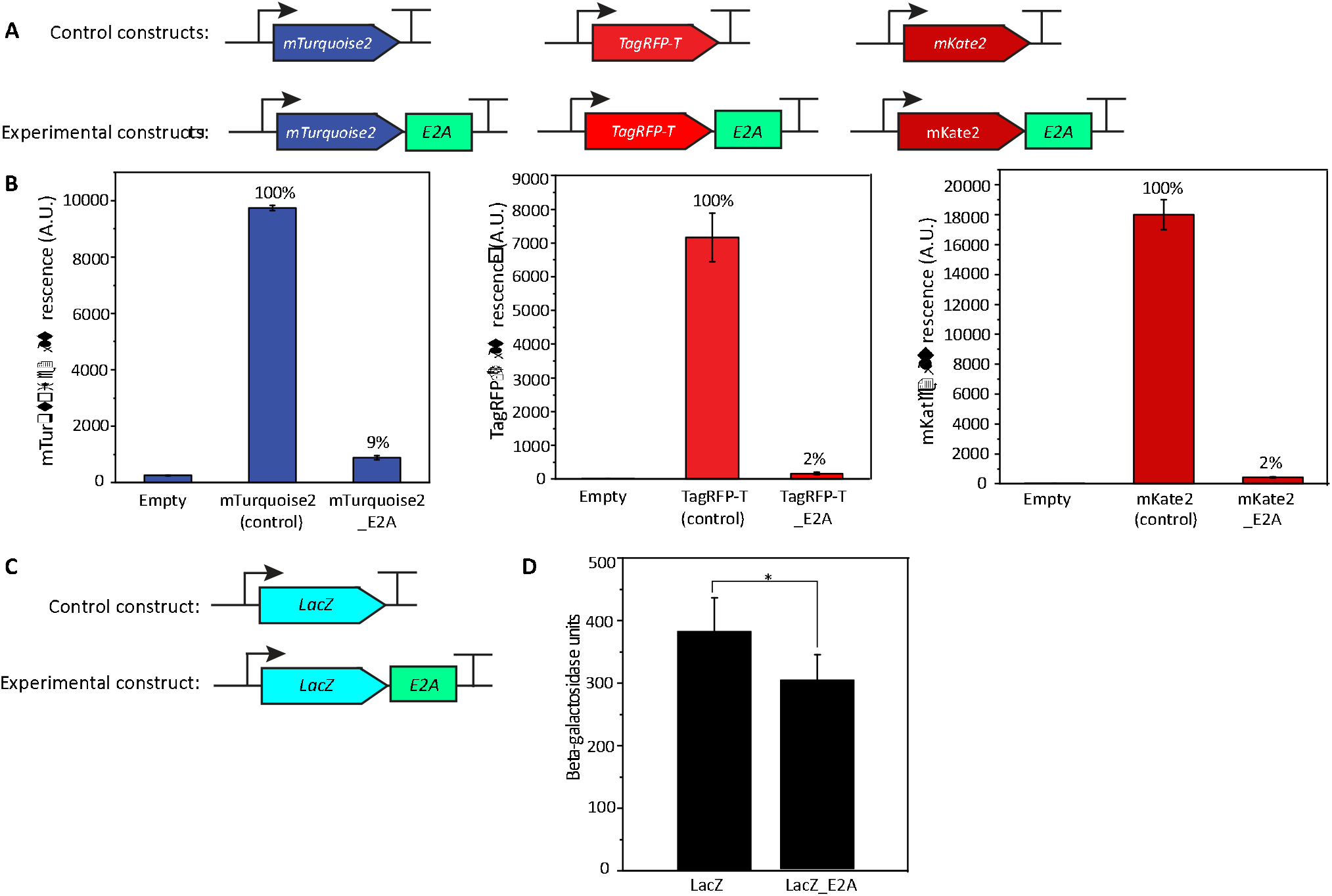
Effect of the C-terminal E2A sequence on protein function. (A) Constructs for testing the effect of the E2A sequence on fluorescent proteins. (B) Normalized fluorescence of mTurquoise2, TagRFP-T, and mKate2 in the empty, control, and experimental constructs. (C) Constructs for testing the effect of the E2A sequence on LacZ protein. (D) The β-galactosidase activities of LacZ with or without the E2A. *: p < 0.1. Data represent the average ± SD of three independent biological replicates.

Since various yeast strains are used for metabolic engineering, we tested the bi-cistronic constructs in four widely used yeast strains, including three haploid strains (BY4741, CEN.PK2-1C, W303) and one diploid strain (CEN.PK2) (Supplementary Figure 8). Fluorescence measurements of both genes were similar across all four strains regardless of which 2A peptide was used, suggesting that the 2A peptide-based polycistronic expression system performs well regardless of the host strain.

### Characterization of the tri-cistronic constructs

To test the 2A peptide polycistronic-expression system beyond expressing two genes, we created tri-cistronic constructs with three fluorescence proteins, TagRFP-T, mTurquoise2, and Venus, linked by two 2A peptide sequences (Figure 4A). We used two different 2A peptides in tri-cistronic constructs to minimize potential homologous recombination. E2A, P2A, and O2A were included because of their high cleavage efficiencies (Figure 2C&F), and a total of six constructs were generated. Overall, the fluorescence of the first protein, TagRFP-T, is the highest, followed by mTurquoise2 and Venus, compared to their monocistronic counterparts (Figure 4B). Fluorescence of TagRFP-T as the first protein had similar relative fluorescence as in the bi-cistronic constructs (56%-80% of the control). The second protein, mTurquoise2, had lower relative fluorescence in the tri-cistronic constructs (41%-63% of control) than the bi-cistronic ones (Figure 2B). This result could be due to the extra C-terminal 2A peptide after mTurquoise2 that negatively affected fluorescence. The third protein Venus had relative fluorescence ranging from 41-52% of its control. Since Venus does not have the C-terminal 2A peptide, the dip in fluorescence is likely due to the increased distance of Venus from the promoter possibly caused by ribosome run-off when the transcript is too long^*46, 47*^. Western blot showed comparable ratios of cleaved Venus to the uncleaved mTurquoise2-Venus except when O2A was the second 2A peptide. This result agreed with the slightly lower cleavage efficiency of O2A in Figure 2F. Thus, 2A peptides can be reliably used to enable tri-cistronic expression in yeast, albeit the expression level of the third gene can be lower than that of the first two.

**Figure 4.**
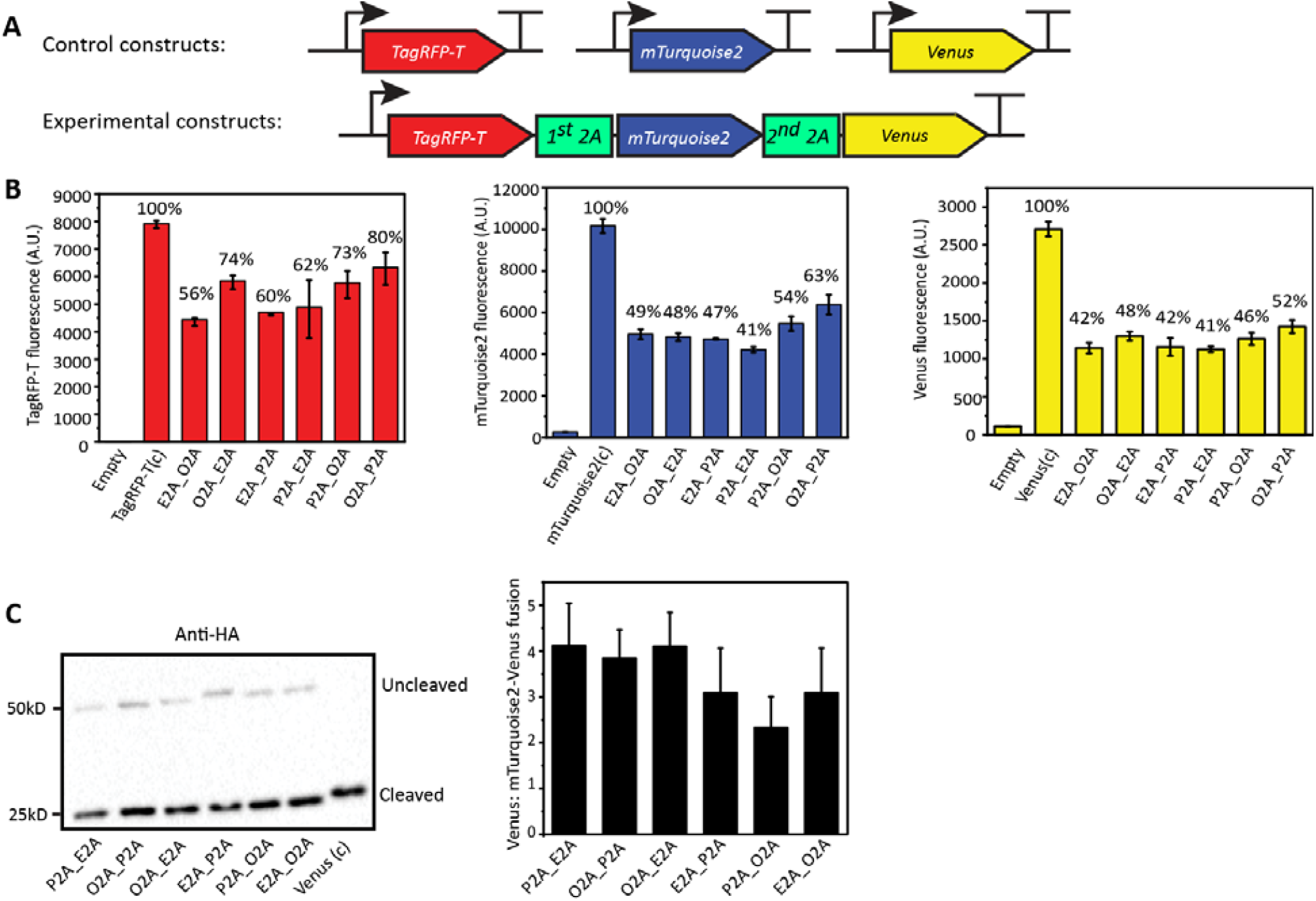
Characterization of tri-cistronic constructs. (A) Control monocistronic and experimental tri-cistronic constructs expressed under the *GAL1* promoter. (B) Normalized fluorescence of TagRFP-T, mTurquoise2, and Venus in the empty, control, and experimental constructs. (C) Western blot to demonstrate the cleavage by the second 2A peptide in the tri-cistronic constructs. Anti-HA primary antibody bound to the HA tag at the C-terminus of Venus. The cleaved Venus was at ∼25kD, and the uncleaved mTurquoise2-2A-Venus fusion protein wa at ∼50kD. The histogram shows the ratio of cleaved Venus to the uncleaved mTurquoise2-2A-Venus fusion protein. Data represent the average ± SD of three independent biological replicates.

### Characterization of the quad-cistronic construct

We extended the 2A peptide-based polycistronic constructs to express four fluorescent protein, mKate2, TagRFP-T, mTurquoise2, and, Venus, linked by three different 2A peptides (Figure 5A). Overall, there was a decrease in relative fluorescence of all four proteins compared with bi- and tri-cistronic constructs (Figure 5B). This observation could result from the increased cellular burden of expressing multiple genes. The decrease of fluorescence in the first three proteins (31%-40% of controls) was likely due to the C-terminal 2A peptides that affected protein function. The last protein, Venus, showed a significantly reduced fluorescence (21% of control), owing to the further increased distance from the promoter. The Western Blot in Figure 5C showed the cleaved Venus protein from the quad-cistronic construct. No uncleaved fusion protein with Venus was detected.

**Figure 5.**
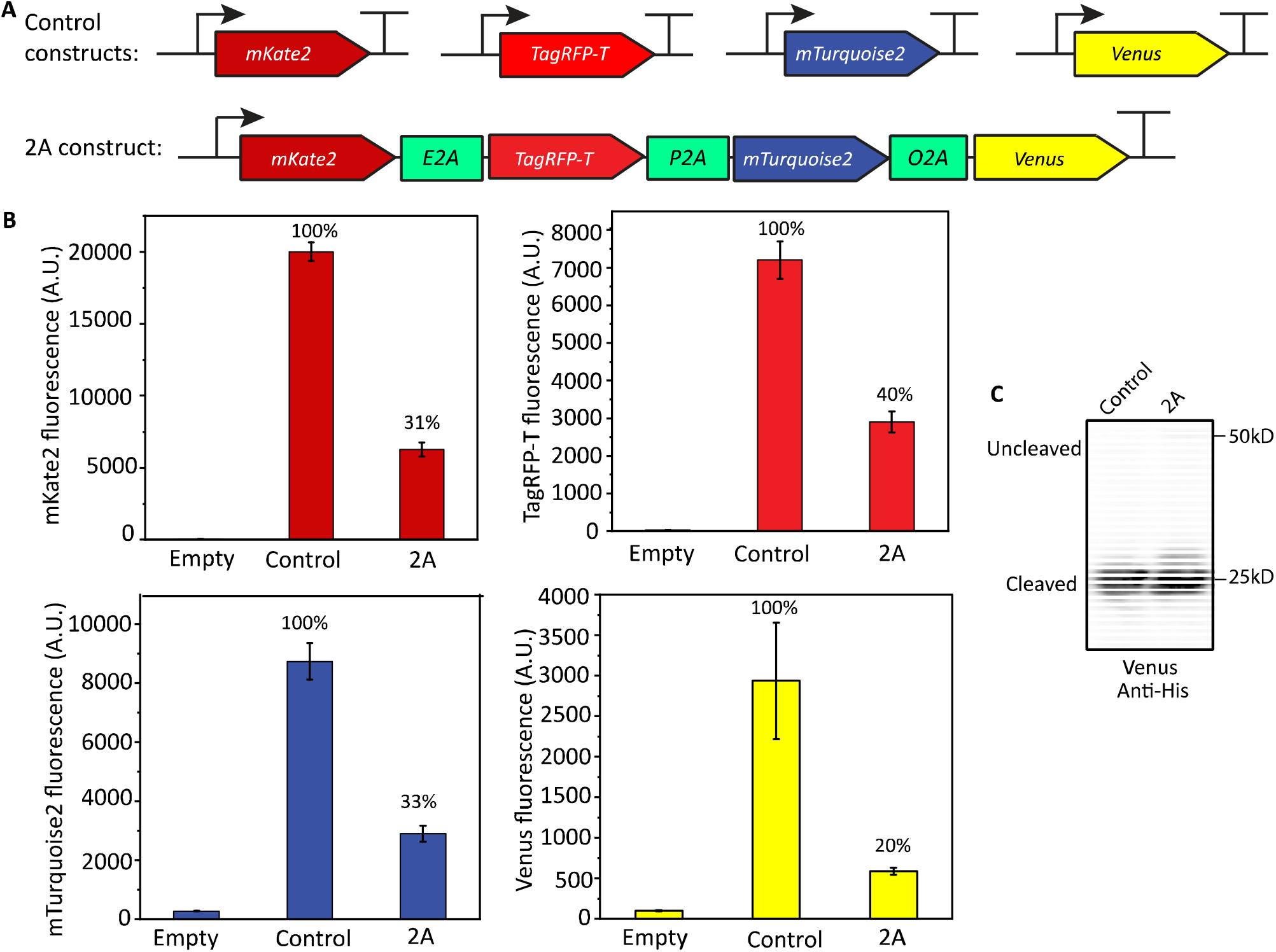
Characterization of quad-cistronic constructs. (A) Control monocistronic and experimental quad-cistronic constructs expressed under the *GAL1* promoter. (B) Normalized fluorescence of mKate2, TagRFP-T, mTurquoise2, and Venus in empty vector, control, and the experimental 2A construct. (C) Western blot showing the cleavage of the third 2A peptide, O2A, in the experimental construct. Anti-his primary antibody detected the 6*His tag at the C-terminus of Venus. The cleaved Venus was at ∼25kD. The uncleaved mTurquoise2-2A-Venus fusion protein was expected at ∼50kD. Data represent the average ± SD of three independent biological replicates.

Thus far, we have characterized the 2A peptide-based polycistronic expression in yeast up to four genes. The positional effect of genes is more pronounced in the quad-cistronic constructs than the tri-cistronic constructs. Although more than four genes can be expressed using this system, we stopped further characterizations due to the limited numbers of fluorescent proteins with non-overlapping light spectra,

### Constructing a metabolic pathway using the 2A peptide-based polycistronic system

Using the 2A peptide-based polycistronic expression system, we constructed a metabolic pathway for producing geraniol in yeast. Geraniol is a valuable mono-terpene extensively used in the fragrance industry^*48*^. To produce geraniol in yeast, we overexpressed *tHMG1* and *IDI1*, the two rate-limiting genes of the mevalonate (MVA) pathway, and the geraniol pyrophosphate (GPP) synthase gene *ERG20*^*ww49*^ and the geraniol synthase gene (*Ocimum basilicum)*, t*Ob*GES^*50*^ (Figure 6A). We assembled four quad-cistronic constructs with four genes in different positions under the *GAL1* promoter. A monocistronic construct (MC) with each gene expressed under the same promoter was used as a control (Figure 6B). Three out of four polycistronic 2A constructs produced similar or higher geraniol compared with the mono-cistronic control, demonstrating that the polycistronic constructs can be used for yeast metabolic engineering (Figure 6C) (Supplementary Figure S9). The C-terminal 2A peptide sequence did not significantly affect the function of each gene. Comparing geraniol titers in strains 2A_1 and 2A_2 indicated that *ERG20*^*ww*^ was the bottleneck of this pathway since the geraniol titer decreased as *ERG20*^*ww*^ was moved farther away from the promoter. The same conclusion could be drawn by comparing strains 2A_3 with 2A_4. Therefore, the positional effects of the 2A polycistronic expression system, combined with the ease of cloning, can be exploited to identify metabolic pathway bottlenecks.

**Figure 6:**
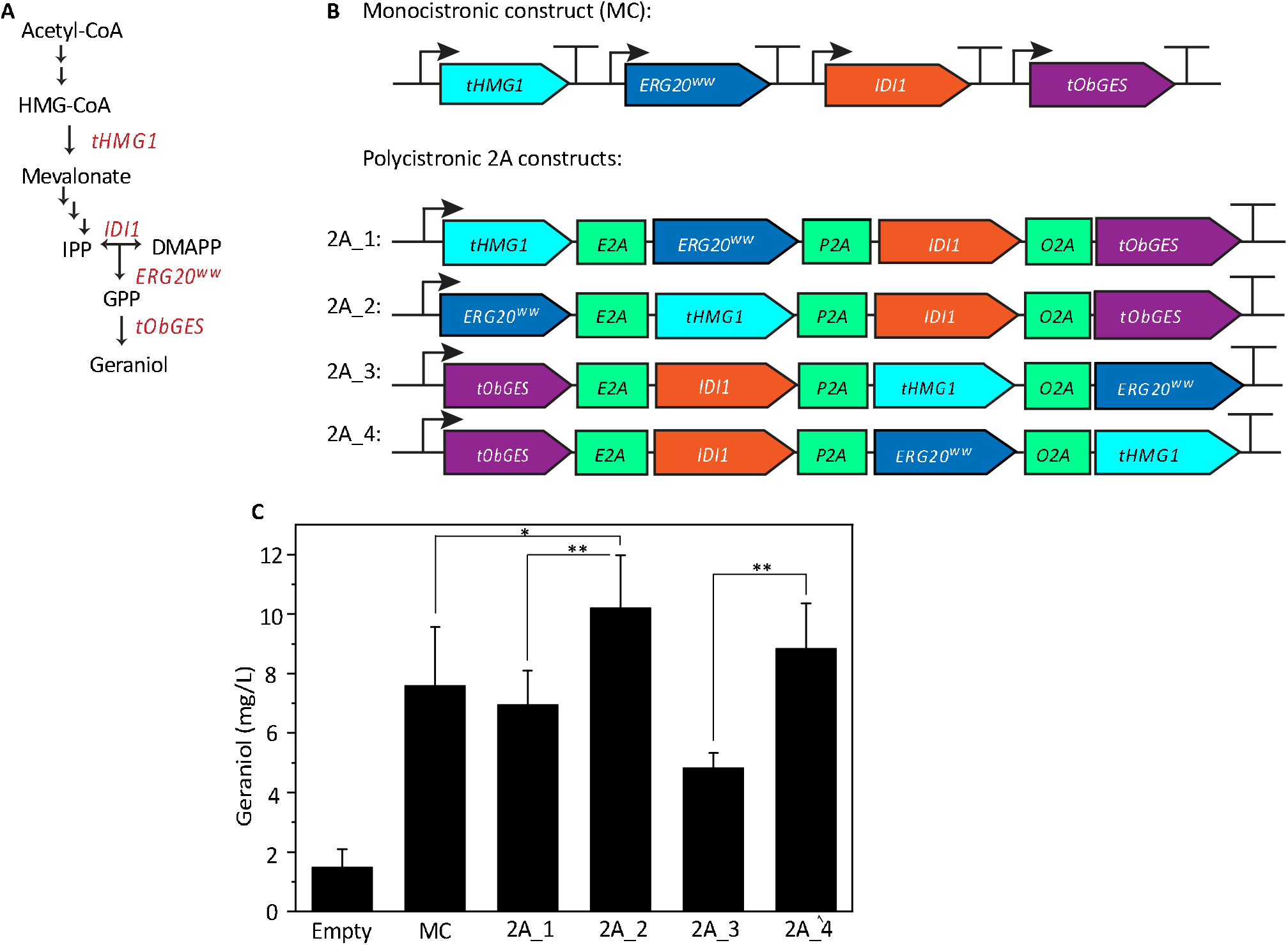
Geraniol production in yeast using the 2A peptide-based quad-cistronic constructs. (A) Mevalonate pathway in yeast for geraniol production. tHMG1: yeast truncated HMG-CoA (β-hydroxy β-methylglutaryl-CoA) reductase without the regulatory domain; IDI1: yeast isopentenyl-diphosphate isomerase; ERG20^*ww*^: a mutant (F96W, N127W) of ERG20 farnesyl pyrophosphate synthetase acting as a geranyl pyrophosphate (GPP) synthase; tObGES: geraniol synthase from Ocimum basilicum. (B) Monocistronic construct (MC) bearing tHMG1, IDI1, ERG20^*ww*^, and tObGES with each under the GAL1 promoter and CYC1 terminator. Polycistronic constructs (2A_1-4) have the above four genes in different positions. (C) Quantification of geraniol produced from yeast transformed with the MC or the 2A_1-4 constructs. *: p < 0.1; **: p < 0.05. Data represent the average ± SD of three independent biological replicates quantified using the geraniol dehydrogenase assay.

### Summary

A polycistronic gene expression system is highly valuable for expressing multiple genes in yeast. To this end, we have developed a 2A peptide-based polycistronic gene expression system compatible with Golden Gate cloning. Such design allowed fast and reliable cloning of multiple genes in the polycistronic expression system. Furthermore, we characterized the bi-, tri-, and quad-cistronic gene expression using fluorescence proteins as reporters. Finally, we constructed a four-gene metabolic pathway to produce geraniol. Geraniol produced from the polycistronic constructs were comparable or superior to the control monocistronic construct.

The performance of this polycistronic expression system is constant across four commonly used yeast strains. Genes farther away from the promoter showed decreased expression in the polycistronic expression system. This phenomenon may be exploited to identify bottlenecks in a metabolic pathway. The C-terminal 2A peptide fused to all except the last protein in the polycistronic system affected protein function in a sequence-dependent manner. We found that the 2A peptide significantly affected fluorescent proteins but had a limited impact on lacZ and the four enzymes for geraniol production. Protease cleavage sequences such as a furin cleavage site may be placed between a CDS and a 2A peptide to eliminate the impact of 2A peptide on protein function^*28, 29*^. Additionally, since the quad-cistronic system developed in this work is at the TU level, and the original yeast MolClo kit allows the cloning of up to six TUs into a multi-gene level plasmid, one can express up to 24 proteins by combining the two systems. Such capacity will undoubtedly facilitate the construction of very long metabolic pathways or elaborated engineering in yeast. Taken together, we expect this work to greatly expedite the wide adaptation of the polycistronic-expressing system for metabolic engineering and other applications in yeast.

## Methods

### Strains and Growth media

The *S. cerevisiae* strain BY4741 (*MATa; his3*Δ*1; leu2*Δ*0; met15*Δ*0; ura3*Δ*0*) was used for gene expression of the mono- and polycistronic constructs. The MVA pathway genes, *tHMG1*, and *IDI1* were amplified from the genome of the strain CEN.PK2-1C (*MATa his3D1 leu2-3,112 ura3-52 trp1-289 MAL2-8C SUC2*). Chemically competent *Escherichia coli* (*E. coli*) DH5α strain was used to propagate plasmids. Transformed *E. coli* cells were selected on the Luria-Bertani plate (LB) with appropriate antibiotics. The wildtype *E. coli* strain MG1655 was used to amplify the *LacZ* gene.

Synthetic dropout media contained 0.67% (w/v) yeast nitrogen base without amino acids (Difco, Franklin Lakes, NJ), 0.1% (w/v) dextrose (Fisher Scientific, Waltham, MA), 2% (w/v) raffinose (Goldbio, St. Louis, MO), 0.07% (w/v) synthetic complete amino acid mix minus histidine (Sunrise Science, Knoxville, TN). Galactose induction media had the same composition, except the 0.1% (w/v) dextrose was replaced with 2% galactose (Fisher Scientific, Waltham, MA). Yeast extract peptone dextrose (YPD) media for preparing yeast competent cells contained 1% (w/v) Bacto™ yeast extract, 2% (w/v) Bacto™ peptone, and 2% (w/v) dextrose.

### Gene synthesis, PCR, and Cloning

The *ERG20*^*WW*^ and t*Ob*GES genes and the 2A peptides were synthesized by IDT (Newark, NJ). PCR amplification was performed using the Phusion High Fidelity DNA Polymerase (NEB, Ipswich, MA) according to the manufacturer’s protocol. All primer sequences were included in Supplementary Table S5. Gibson assembly^*51*^ was used to insert the *LacZ* and the *LacZ_E2A* into the pCEN-HIS vector. Golden Gate assembly was performed for all the other constructs as outlined in Mukherjee et al., 2021^*52*^. In brief, each reaction mix contained: 1 μl of equimolar concentration (∼20fmol) of each DNA fragment or plasmid, 1 μl of T4 DNA ligase buffer (NEB, Ipswich, MA), 0.5μl of restriction enzyme (BsaI-HFv2 or Esp3I), 0.5 μl of T4 ligase (NEB, Ipswich, MA), followed by adjusting the volume to 10 μl. Reaction mixes were incubated in a thermocycler using the following program: 25-35 cycles of digestion and ligation at 37 °C for 5 min, and 16 °C for 5 min respectively, followed by a final digestion step at 50 °C for 10 min, and a heat inactivation step at 80 °C for 10 min. The final digestion and heat inactivation steps were omitted for assembling the intermediate TU vector^*14, 52*^. The sequences of all part plasmids and pCEN-His plasmids were confirmed using Sanger sequencing (GeneWiz, South Plainfield, NJ).

### Measuring the fluorescence of the polycistronic constructs

Yeast competent cells were transformed with polycistronic constructs containing fluorescent proteins using the Frozen-EZ yeast transformation II kit (Zymo Research, Irvine, CA) according to the manufacturer’s protocol. The transformation mixture was plated on histidine dropout (SD-His) plates and incubated at 30 °C for two days. Colonies were picked and grown overnight in 5 ml liquid dropout media at 30 °C with shaking at 200 rpm. The overnight culture was inoculated at an initial OD_600_ of 0.1 into the induction media containing galactose and grown at 30 °C with shaking at 200 rpm for 24 hours. 200 μl of the culture was transferred into a 96-well plate. Fluorescence was measured using a Tecan (Morrisville, NC) Spark microplate reader. Excitation and emission wavelengths used were: mTurquoise2 at 435 nm / 478 nm, TagRFP-T at 555nm / 585nm, Venus at 485nm / 530nm, mKate2 at 588nm / 633nm, respectively. The fluorescence readings were normalized to the OD_600_.

### β-galactosidase assay

The β-galactosidase assay was performed according to the protocol described in the Clontech (Mountain View, CA) Yeast Protocol Handbook. In brief, yeast transformed with plasmids bearing the *LacZ* gene was grown as described in the paragraph above. 1.5 ml of the induced culture was collected at OD_600_ of 0.8 and pelleted at 10,000 x g for 1 min, then resuspended in 1.5 ml of Z buffer (16.1 g/l Na_2_HPO_4_*7H_2_O, 5.5 g/l NaH_2_PO_4_*H_2_O, 0.75 g/l KCl, 0.246 g/l MgSO_4_*7H_2_O; pH 7.0). Cells were pelleted again and resuspended in 300 μl of Z buffer (concentration factor: 5). Next, 100 μl of cell suspension was frozen in liquid nitrogen and thawed at 37 °C for 1 min. A total of three freeze-and-thaw cycles were performed to lyse the cells. To initiate the assay, 700 μl of Z buffer, 0.27% 2-mercaptoethanol, and 160 μl of 4 mg/ml ONPG (*ortho*-nitrophenyl-β-galactoside in Z buffer) were added to the cell lysate. The assay mixtures were incubated at 30 °C for ∼15 min until the yellow color developed, followed by the addition of 400 μl of 1 M Na_2_CO_3_. Cells were pelleted at 10,000 x g for 10 min, and the supernatant was used to measure the absorbance at 420 nm using a UV-vis spectrophotometer (Thermo Fisher Scientific, Waltham, MA). β-galactosidase units were calculated according to the equation:

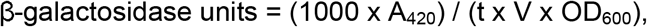

where t is the time to develop the yellow color and V is 0.1 x concentration factor.

### Yeast protein extraction

Cell pellet from 1.5 ml of culture was resuspended in 250 μl of 1% (v/v) 2-mercaptoethanol and 0.25 M NaOH. After 10 min on ice, 160 μl of 50% trichloroacetic acid (TCA) was added to the cell suspension, then incubated for 10 min on ice. After centrifugation at 16,000 x g for 10 min, the supernatant was removed, and the pellet was allowed to air dry. The pellet was then resuspended in 200 μl sample buffer containing 2% (w/v) 2-mercaptoethanol, 125 mM Tris-HCl, 20% (w/v) glycerol, 4% (w/v) sodium dodecyl sulfate (SDS) and 4% (w/v) bromophenol blue. This sample buffer was used for protein gel electrophoresis and Western blot.

### Western blot

Western Blot was performed with modifications from the protocol described in Wang et al., 2012^*53*^. In brief, 20 μg protein from each sample was separated by SDS-PAGE (polyacrylamide gel electrophoresis), then transferred to the polyvinylidene fluoride (PVDF) membrane (Bio-Rad, Hercules, CA), which was then blocked with 5% (w/v) non-fat milk in TBST buffer (20 mM Tris, 150 mM NaCl, 0.1% Tween-20) at room temperature for three hours. Primary antibody (Anti-His; Invitrogen, Waltham, MA or Anti-HA; Sigma Aldrich, St.Louis, MO) was added to the blocking solution at 1:2,000 dilution and incubated overnight at 4 °C. The PVDF membrane was then washed 3 × 10 min with TBST and incubated with horseradish peroxidase (HRP)-conjugated goat anti-mouse (diluted 1:20,000; Bio-Rad, Hercules, CA) secondary antibody for one hour at room temperature followed by 6 × 10 min washes with TBST and detection using the Radiance Plus chemiluminescence substrate (Azure Biosystems, Dublin, CA) and ChemiDoc XRS+ image analyzer (Bio-Rad, Hercules, CA). The band intensities were quantified using the ImageJ software. The following formula was used to measure the cleavage efficiency^*41*^:

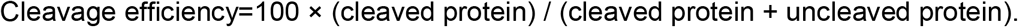

### Geraniol production and quantification

Yeast colonies transformed with MC and 2A_1-4 plasmids were grown overnight in 5 ml dropout media at 30 °C with shaking at 200 rpm. The overnight culture was inoculated at an initial OD_600_ of 0.1 into the induction media and grown at 30 °C with shaking at 200 rpm for 24 hours. 1 ml of the culture was pelleted at 16,000 x g for 1 min, and the supernatant was used to quantify geraniol using the geraniol dehydrogenase (Gedh) assay^*54*^. In brief, 50 μl of the supernatant was mixed with 50 μl of the reaction mix containing 10 μl of 200 mM Tris-HCl (pH 8.0), 4 μl of 4 mM NAD^+^ (Goldbio, St. Louis, MO), 4 μl of 4 mM resazurin sodium salt (Acros Organics, Belgium), 0.002 U purified *Castellaniella defragrans* geraniol dehydrogenase, and 1 μl of 100 U diaphorase (Sigma Aldrich, St.Louis, MO). For preparing the geraniol standard curve, 10X of each geraniol concentration was prepared by dissolving authentic geraniol standard (Acros Organics, Belgium) in acetone. Next, 5 μl of the 10X geraniol and 45 μl of double distilled H_2_O was added to the 50 μl reaction mix, such that the final geraniol concentration is 1X. The geraniol standard curve is in Supplementary Figure 10. Each reaction was incubated at room temperature for 45 min, and fluorescence was recorded at the excitation and emission of 530 nm and 590 nm, respectively, using a Tecan Spark microplate reader (Morrisville, NC).

## Supporting Information

The supplementary information includes supporting instructions (figures, tables, and text) on how to design primers and assemble the poly-cistronic constructs; supporting figures to show the effect of longer linkers on 2A peptide function, the effectiveness of four 2A peptides in four yeast strains; supporting chromatograms and MS spectra showing the authentic standard and geraniol produced by the quad-cistronic construct; supporting tables that list of plasmids and primers used in the study.

## Acknowledgment

The authors are grateful to Dr. Sarah Walker for providing access to the Tecan Spark microplate reader and Dr. Valerie Frerichs from the Chemistry Instrument Center (CIC) at University at Buffalo for the mass spectrometry work. We also thank Dr. Laura Rusche for providing the yeast haploid strain W303 (LRY 1006) for this study. This project was supported by the Research Foundation for the State University of New York (71272) to Z. Q. Wang and the National Science Foundation (CHE-1919594) to the CIC.

