## Supplemental information for "A Well-Characterized Polycistronic-Like Gene Expression System in Yeast"

**Supplementary materials and methods**

***S. cerevisiae* strains***.* Four strains were used for screening four 2A peptides: BY4741 (*MATa his3Δ1 leu2Δ0 met15Δ0 ura3Δ0*), CEN.PK2-1C (*MATa his3Δ1 leu2-3, 112 ura3-52 trp1-289 MAL2-8C SUC2*), W303 (*MATa{leu2-3,112 trp1-1 can1-100 ura3-1 ade2-1 his3-11,15}[phi^+^]*), and

CEN.PK2 (*MATa/α his3Δ1/his3Δ1 leu2-3,112/leu2-3,112 ura3-52/ura3-52 trp1-289/trp1-289 MAL2-8C/MAL2-8C SUC2/SUC2*).

**GC-MS for geraniol detection**. For geraniol extraction from the engineered yeast, 1 ml culture was centrifuged at 16,000 x rpm for 1 min. 500 μl of the supernatant was mixed with 500 μl hexane and vortexed at the highest speed for 1 min, then centrifuged at 16,000 x rpm for 2 mins. 500 μl of the hexane layer was collected and used for GC-MS. Geraniol concentration of 25 mg/L was prepared by dissolving authentic geraniol standard in hexane and directly used for GC-MS. Geraniol was detected using Thermo Trace 1300 Gas Chromatograph and Thermo Q-exactive Orbitrap Mass Spectrometer (Waltham, MA). 5 μL of the sample was injected into a Thermo Scientific TraceGOLD TG-5SILMS column (30 m long, 0.25 mm inner diameter, 0.25 μm film thickness) using helium as the carrier gas (1 ml/min). The injector was held at 200 °C. The oven was held at 40 °C for 4 mins, followed by ramping up to 280 °C at a rate of 20 °C/min and then holding at 280 °C for 2 mins. The MS transfer line was at 250 °C, and the source temperature was 200 °C.  The resolution was set to 60,000.  The mass range monitored was 39 – 200 M/Z in the positive ion mode. The MS was set to monitor total ion counts. Geraniol eluted at 10.24 mins.

**Cloning 2A peptide-based multigene plasmids.**

*Primer design for modular Golden Gate cloning:*

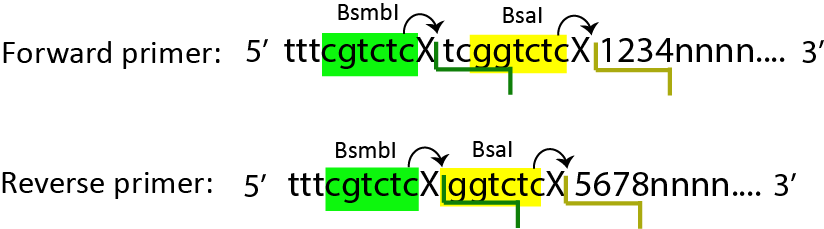

**Figure S1:** Design of forward and reverse primers.

Both the forward and reverse primers have the BsmbI recognition (cgtctc) and cut site (tcgg and ggtc for forward and reverse primers respectively), and BsaI recognition (ggtctc) and cut sites that include four nucleotide overhangs specific to the DNA part*^1^*.

*Designing the Golden-Gate overhangs for cloning 2A polycistronic constructs:*

All 2A peptides have an N-terminal GSG linker and the conserved C-terminal PGP sequence. We took advantage of the wobble position of the amino acid codons to create unique four-nucleotide overhangs for the 2A peptide polycistronic constructs, as detailed below.

*Assembly of the bi-cistronic constructs:*

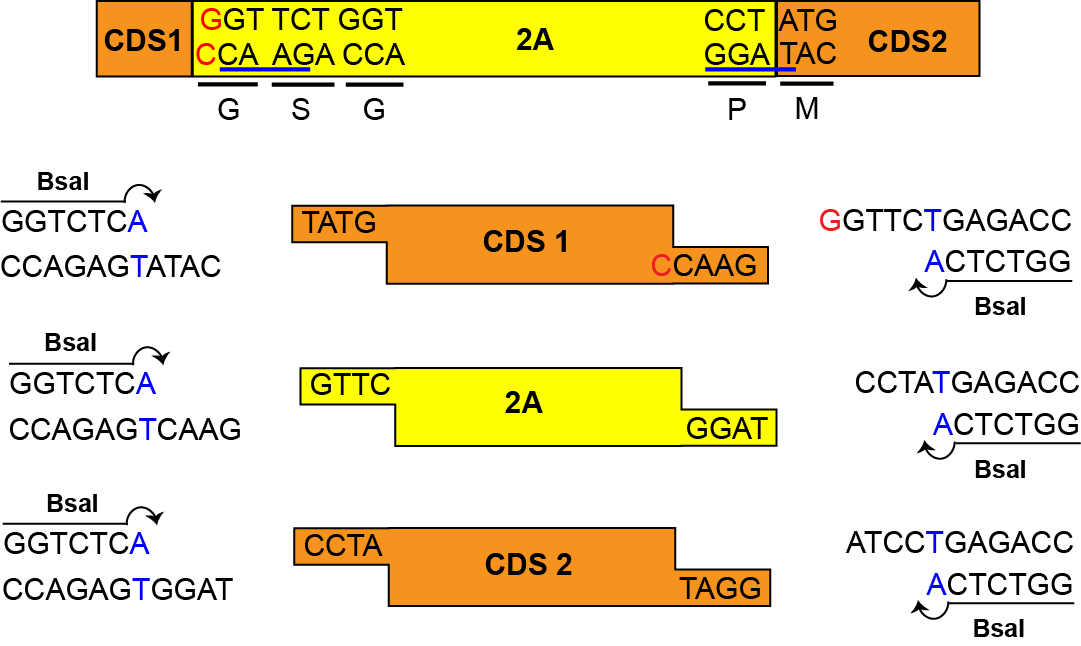

**Figure S2:** The four-nucleotide overhangs for bi-cistronic constructs. Letters in red are the additional nucleotides added to complete the glycine (GGT) in the GSG linker.

**Table S1:** Primer design for bi-cistronic constructs. Nucleotides in bold are BsmbI and BsaI recognition sites. Nucleotides in capital *italics* are the part-specific overhangs to be added to the primers. ‘*nnnn*…’ indicate nucleotides that are specific to the DNA template. Letters in red are the additional nucleotides added to complete the glycine (GGT) in the GSG linker.

| **DNA part** | **Forward Primer (5’-3’)** | **Reverse Primer (5’-3’)** |
| --- | --- | --- |
| CDS 1 | ttt**CGTCTC**gTC**GGTCTC**a*TATGnnnn…* | ttt**CGTCTC**g**GGTCTC**a*GAACCnnnn…* |
| 2A | ttt**CGTCTC**gTC**GGTCTC**a*GTTCnnnn…* | ttt**CGTCTC**g**GGTCTC**a*TAGG*n*nnn…* |
| CDS 2 | ttt**CGTCTC**gTC**GGTCTC**a*CCTAnnnn…* | ttt**CGTCTC**g**GGTCTC**a*GGATnnnn…* |

The CDS1 is flanked by a 5’ TATG overhang to connect with the promoter and a 3’ GTTC to connect with the 2A peptide. TATG includes the start codon ATG, so the coding sequence should begin from the second codon*^1^*. The 3’ GTTC of CDS1 is derived from the glycine (G**GT**) and serine (**TC**T) of the 2A peptide. An extra ‘C’, complementary to ‘G’, needs to be added to the reverse primer before the overhang to complete the glycine (GGT) sequence. The 2A peptide is flanked by the 5’ GGTC and 3’ CCTA. CCTA is derived from the proline (**CCT**) codon and the first codon of the second CDS (**A**TG). The CDS2 is flanked by the 5’ CCTA and the 3’ ATCC which connects to a terminator*^1^*.

*Assembly of the tri-cistronic constructs:*

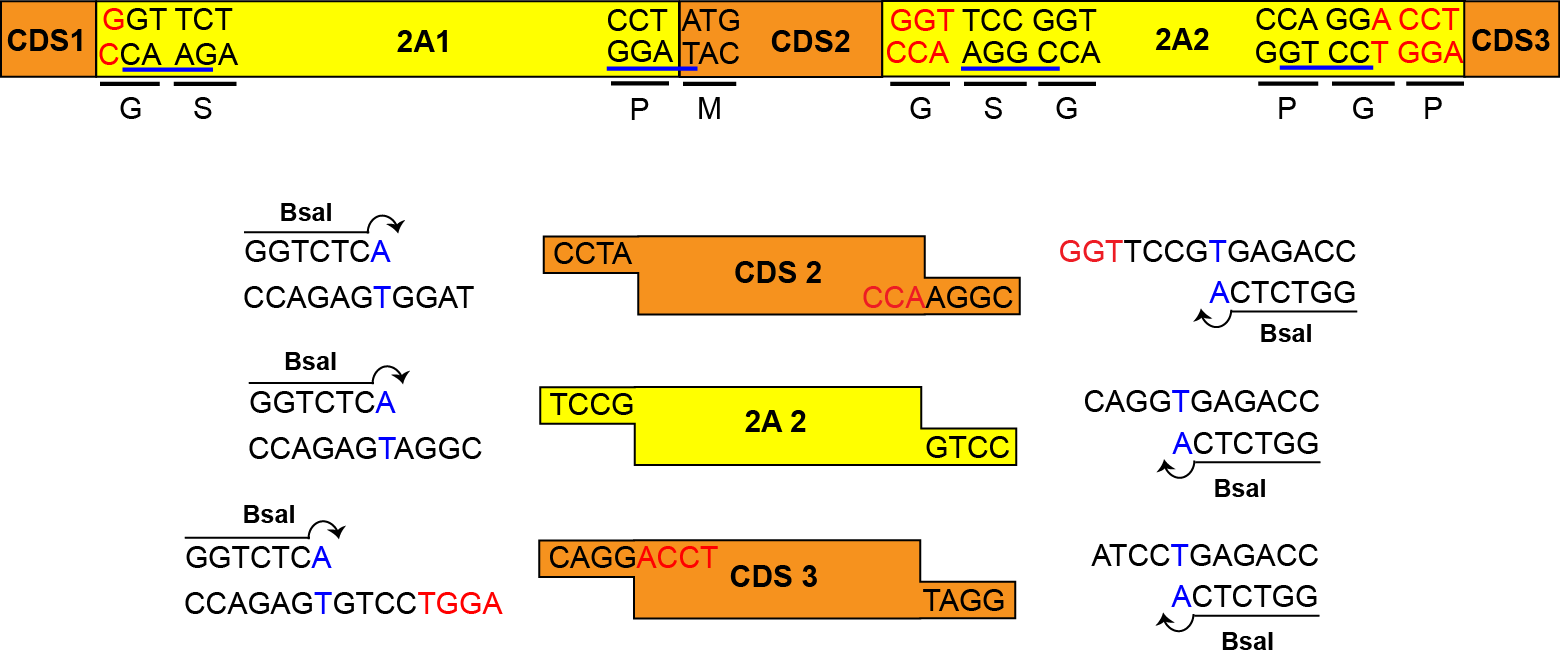

**Figure S3:** The four-nucleotide overhangs for tri-cistronic constructs. Letters in red are the additional nucleotides added to complete the second 2A peptide.

**Table S2:** Primer design for tri-cistronic constructs. Nucleotides in bold are BsmbI and BsaI recognition sites. Nucleotides in capital *italics* are the part-specific overhangs to be added to the primers. ‘*nnnn*…’ indicate nucleotides that are specific to the DNA template. Letters in red are the additional nucleotides added to complete the second 2A peptide.

| **DNA part** | **Forward Primer (5’-3’)** | **Reverse Primer (5’-3’)** |
| --- | --- | --- |
| CDS 1 | ttt**CGTCTC**gTC**GGTCTC**a*TATGnnnn…* | ttt**CGTCTC**g**GGTCTC**a*GAACCnnnn…* |
| 2A 1 | ttt**CGTCTC**gTC**GGTCTC**a*GTTCnnnn…* | ttt**CGTCTC**g**GGTCTC**a*TAGGnnnn…* |
| CDS 2 | ttt**CGTCTC**gTC**GGTCTC**a*CCTAnnnn…* | ttt**CGTCTC**g**GGTCTC**aC*GGAACCnnnn…* |
| 2A 2 | ttt**CGTCTC**gTC**GGTCTC**a*TCCGnnnn…* | ttt**CGTCTC**g**GGTCTC**a*CCTGnnnn…* |
| CDS 3 | ttt**CGTCTC**gTC**GGTCTC**a*CAGGACCTnnnn…* | ttt**CGTCTC**g**GGTCTC**a*GGATnnnn…* |

The four-nucleotide overhangs for CDS1, the first 2A peptide, and the 5’ of the CDS2 in a tri-cistronic construct are identical to that of the bi-cistronic constructs (**Figure S2**). The 3’ overhang for the CDS2 is TCCG, derived from the serine (**TCC**) and the 2^nd^ glycine (**G**GT) of the GSG linker in front of the 2^nd^ 2A peptide. Thus, an additional GGT is needed before the TCCG to encode the 1^st^ glycine in the linker (**Figure S3** in red). The 2^nd^ 2A peptide is flanked by the 5’ TCCG and the 3’ CAGG, respectively. The CAGG is derived from the codons of the 1^st^ proline (C**CA**) and the glycine (**GG**A) of the conserved 3’ PGP sequence in 2A peptides. The CDS3 is flanked by the 5’ CAGG and the 3’ ATCC, respectively. To complete the 3’ PGP in the 2^nd^ 2A peptide, the 5’ of CDS3 also includes the A of glycine (GG**A**) and **CCT** encoding the last proline (**Figure S3** ACCT in red). The 3’ ATCC of CDS3 connects to a terminator*^1^*.

*Assembly of the quad-cistronic constructs:*

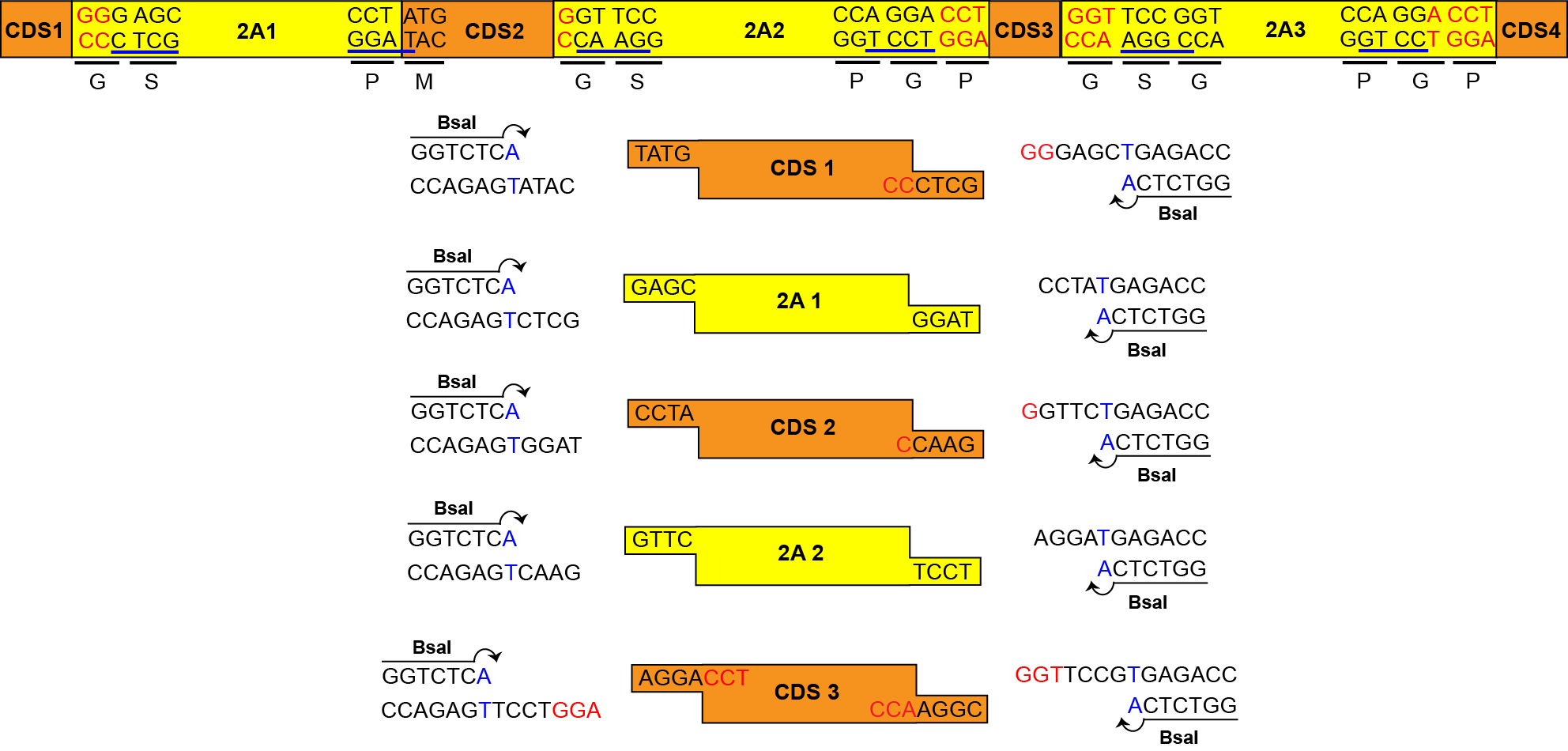

**Figure S4:** The four-nucleotide overhangs for quad-cistronic constructs. Letters in red are the additional nucleotides to complete 2A peptides.

**Table S3:** Primer design for quad-cistronic constructs. Nucleotides in bold are BsmbI and BsaI recognition sites. Nucleotides in capital *italics* are the part-specific overhangs to be added to the primers. ‘*nnnn*…’ indicate nucleotides that are specific to the DNA template. Letters in red are the additional nucleotides added to complete the second 2A peptide.

| **DNA part** | **Forward Primer (5’-3’)** | **Reverse Primer (5’-3’)** |
| --- | --- | --- |
| CDS 1 | ttt**CGTCTC**gTC**GGTCTC**a*TATGnnnn…* | ttt**CGTCTC**g**GGTCTC**a*GCTCCCnnnn…* |
| 2A 1 | ttt**CGTCTC**gTC**GGTCTC**a*GAGCnnnn…* | ttt**CGTCTC**g**GGTCTC**a*TAGGnnnn…* |
| CDS 2 | ttt**CGTCTC**gTC**GGTCTC**a*CCTAnnnn…* | ttt**CGTCTC**g**GGTCTC**a*GAACCnnnn…* |
| 2A 2 | ttt**CGTCTC**gTC**GGTCTC**a*GTTCnnnn…* | ttt**CGTCTC**g**GGTCTC**a*TCCTnnnn…* |
| CDS 3 | ttt**CGTCTC**gTC**GGTCTC**a*AGGACCTnnnn…* | ttt**CGTCTC**g**GGTCTC**a*CGGAACCnnnn…* |
| 2A 3 | ttt**CGTCTC**gTC**GGTCTC**a*TCCGnnnn…* | ttt**CGTCTC**g**GGTCTC**a*CCTGnnnn…* |
| CDS 4 | ttt**CGTCTC**gTC**GGTCTC**a*CAGGACCTnnnn…* | ttt**CGTCTC**g**GGTCTC**a*GGATnnnn…* |

CDS1 is flanked by TATG at the 5’ and GAGC at the 3’ end. GAGC is derived from the glycine (GG**G**) and serine (**AGC**) of the GSG linker. Thus, the reverse primer for CDS1 also needs two extra CC to complete the 1^st^ glycine in the GSG linker. The 1^st^ 2A peptide is flanked by GAGC and CCTA. CCTA derives from codons for the C-terminal proline (**CCT**) and the N-terminal methionine (**A**TG) of CDS2. The 2^nd^ CDS is flanked by the 5’ CCTA and the 3’ GTTC. GTTC derives from the codons of the 1^st^ glycine (G**GT**) and the serine (**TC**C) in the GSG linker of the 2^nd^ 2A peptide. Thus, the reverse primer of the 2^nd^ CDS needs an extra C to complete the codon for the 1^st^ glycine. The 2^nd^ 2A peptide is flanked by the 5’ GTTC and the 3’ AGGA. AGGA is derived from the 1^st^ proline (CC**A**) and the next glycine (**GGA**) of the C-terminus PGP sequence in the 2A peptide. The 3^rd^ CDS is flanked by the 5’ AGGA and 3’ TCCG. For the 5’ end, the codon for the terminal proline (CCT) of the 2^nd^ 2A peptide needs to be included in the forward primer in addition to the AGGA overhang. The 3’ overhang TCCG derives from the codons for the serine (**TCC**) and the 2^nd^ glycine (**G**GT) of the GSG linker in front of the 3^rd^ 2A peptide. Thus, the anti-codon of the 1^st^ glycine (GGT), ACC, should be included in the reverse primer of CDS3. The 3^rd^ 2A peptide and the 4^th^ CDS have the same overhangs as the 2^nd^ 2A peptide and the 3^rd^ CDS of the tri-cistronic constructs (**Figure S3**).

*Cloning DNA parts into entry vectors:*

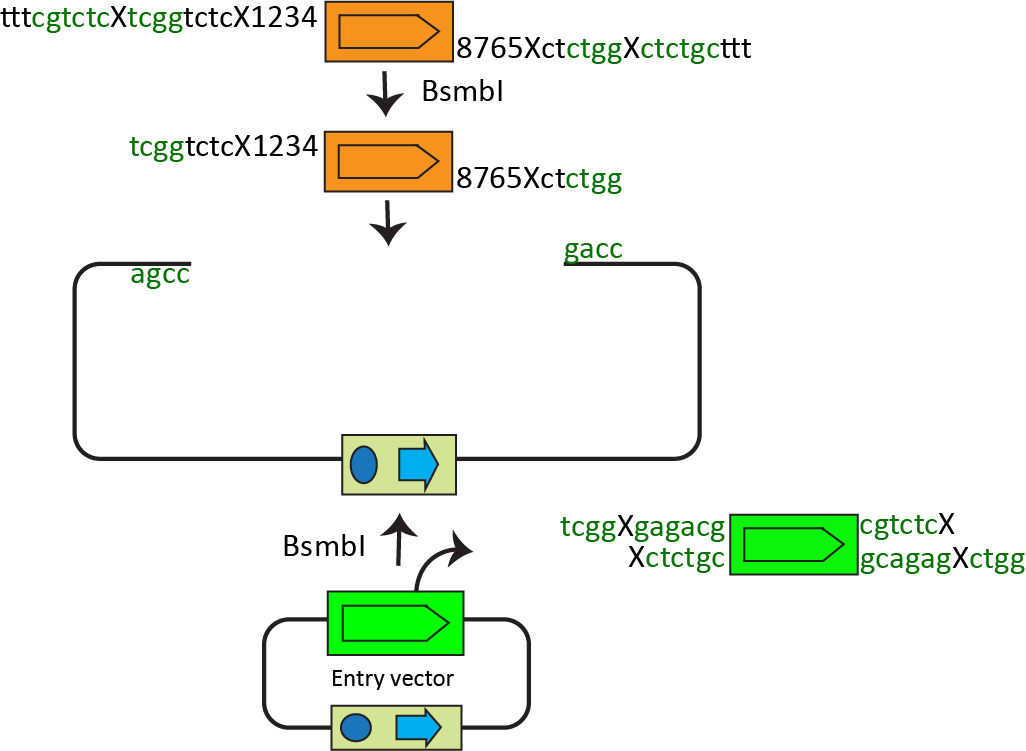

**Figure S5**: Cloning of a part plasmid.

The complete protocol for cloning a part plasmid can be found in Mukherjee, et. al.*^2^*. Briefly, the BsmbI-digested DNA part and the cut entry vector ligate in a Golden-Gate reaction to generate the part plasmid, which retains the two BsaI cutting sites flanking the DNA part.

*Assembly of polycistronic transcription units (TUs):*

The polycistronic TU contains the following DNA parts: left and right connectors, a promoter, CDSs (2-4), 2A peptides (1-3), a terminator, a yeast origin of replication, a yeast selection marker, an *E. coli* selection marker plus an origin of replication, and a yeast origin of replication. Except for the CDSs and the 2A peptides, all the other parts are available in the MoClo-YTK (Addgene kit#1000000061, depositing lab: John Deuber). The 2A peptide plasmids will be deposited to Addgene. The most straightforward way to assemble the TUs is to assemble all the individual parts in one reaction using the BsaI enzyme. However, the efficiency is very low.

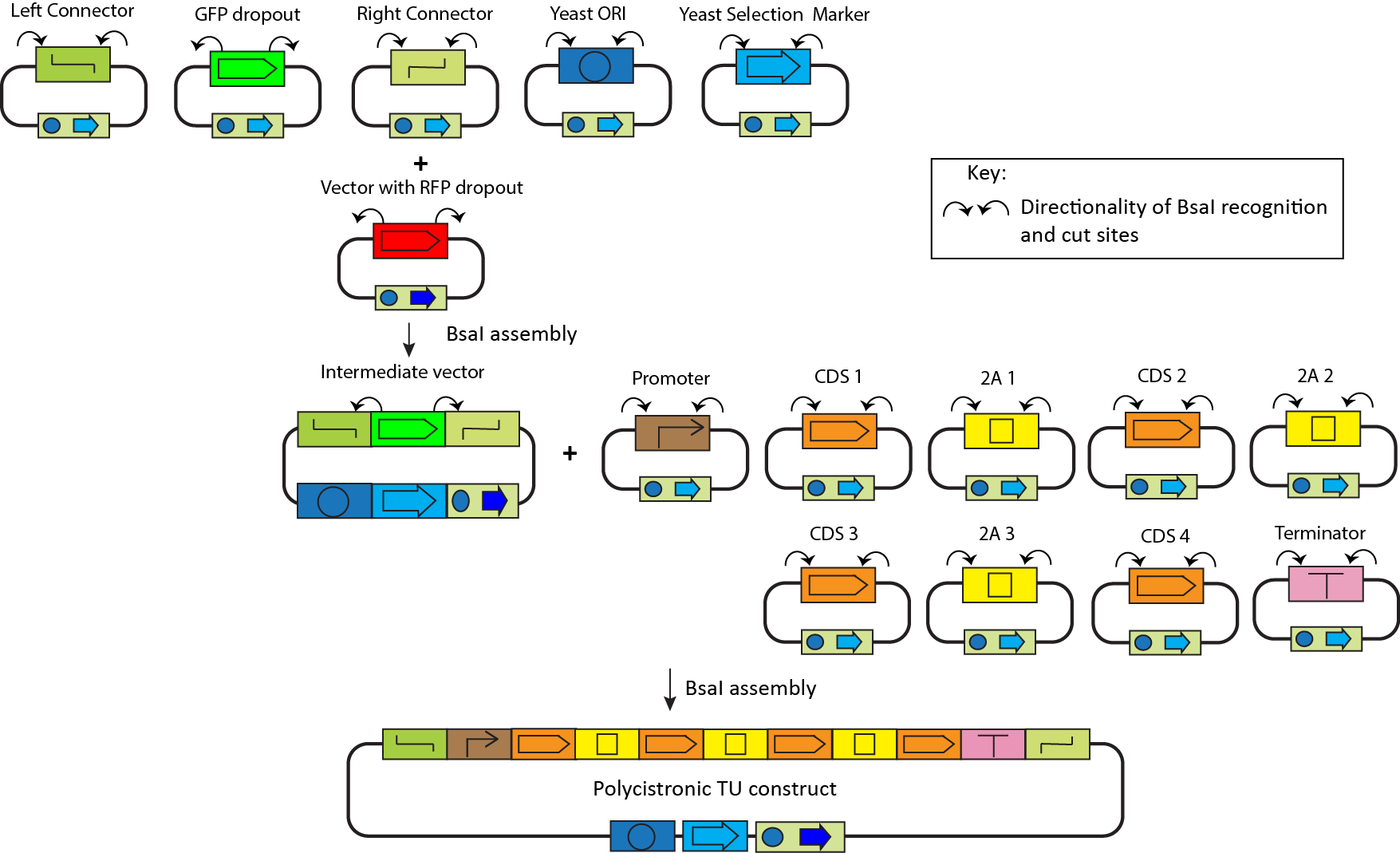

**Figure S6:** Assembly of a 2A peptide-based polycistronic transcription unit.

An alternative and more practical approach is to clone an intermediate vector including the common parts and a GFP dropout with the left and right connectors. Then, clone the promoter, CDSs, 2A peptide(s), and the terminator into the intermediate vector (**Figure S6**). This strategy makes cloning easy by reducing the number of parts *^1, 2^*. If one does not want to vary the promoter and terminator in a set of 2A polycistronic vectors, then one may assemble an intermediate vector that already has the promoter and the terminator flanking the GFP dropout. This will further reduce the number of parts to be assembled. We used this strategy to clone the tri- and quad-cistronic constructs in this work.

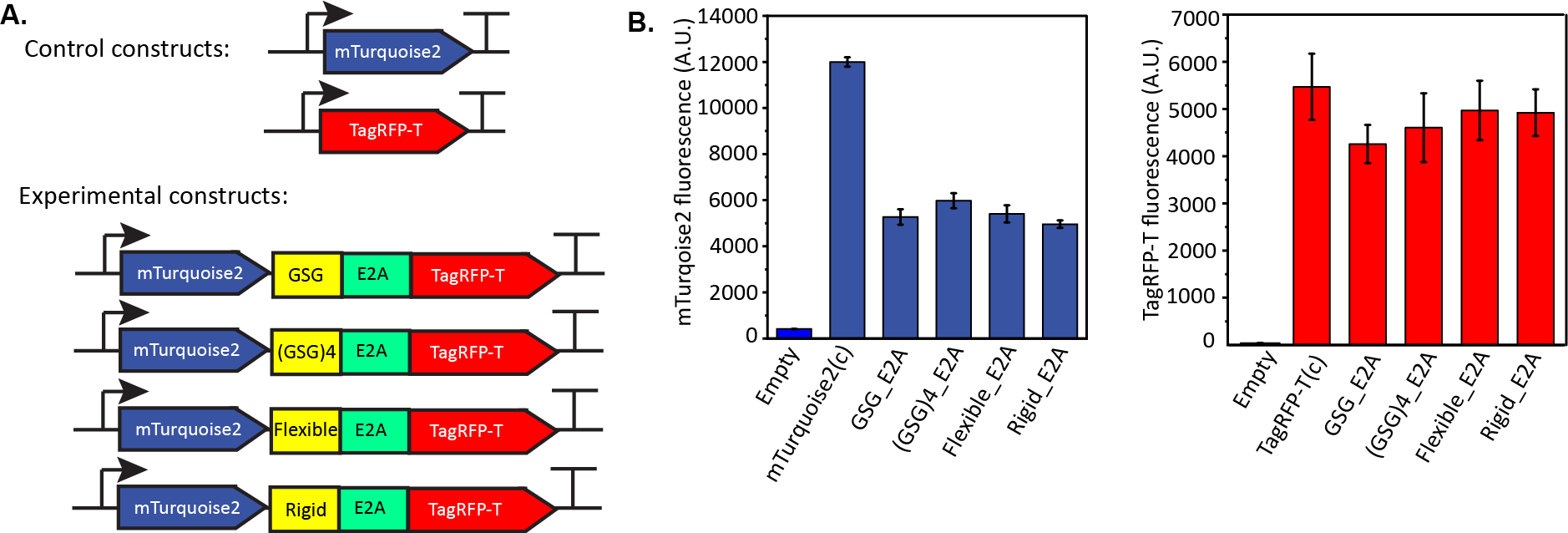

**Figure S7:** Effects of various linker peptides on 2A peptide efficiency. (A) Control monocistronic and experimental bi-cistronic constructs with E2A preceded by four linker peptides, GSG*^3^*, (GSG)4, Flexible, and Rigid*^4^*, respectively. (B) Normalized fluorescence of mTurquoise2 and TagRFP-T in the empty vector, control, and experimental constructs. Data represent the average ± SD of three independent biological replicates.

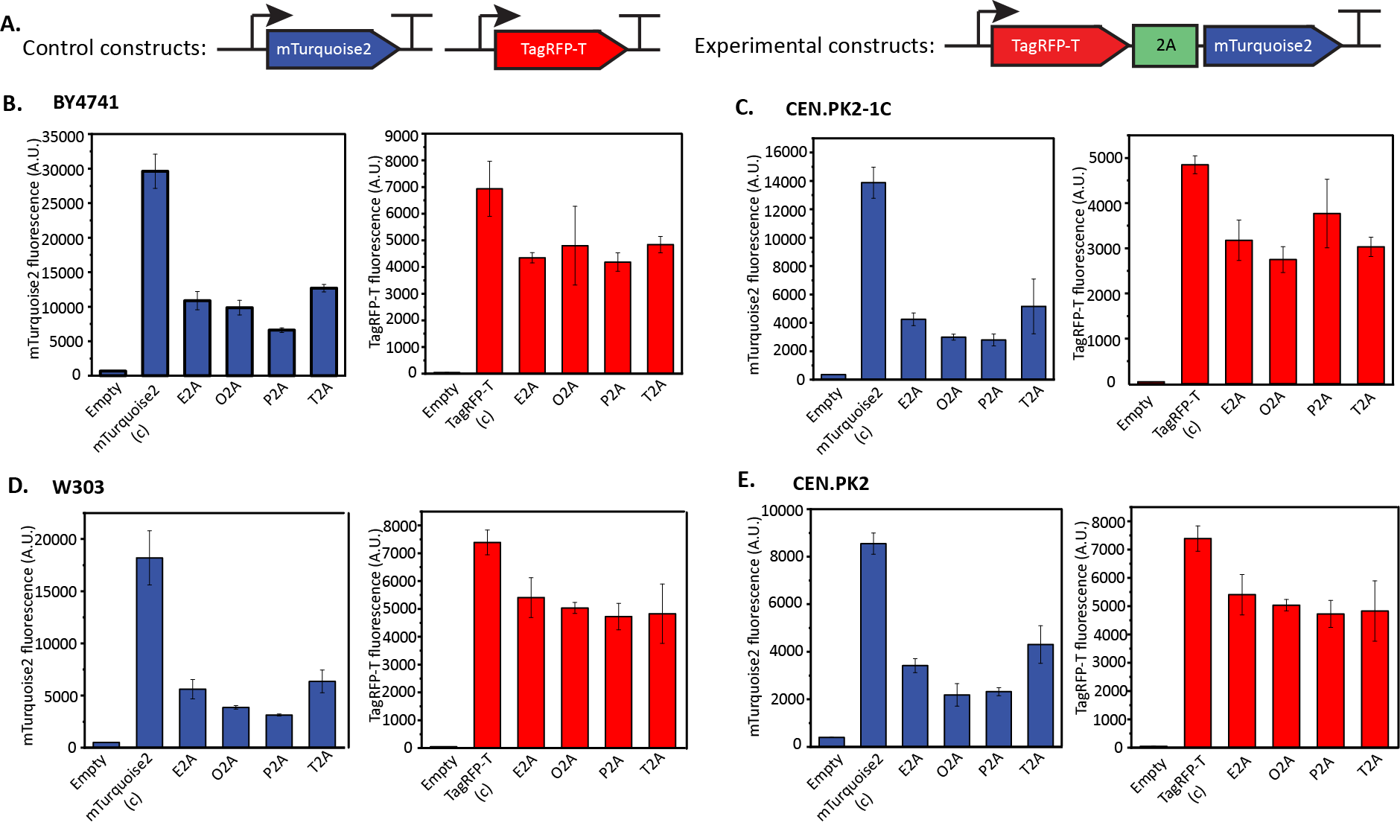

**Figure S8:** Screening four 2A peptides (E2A, O2A, P2A, and T2A) in four yeast strains. (A) Control monocistronic and experimental bi-cistronic constructs. Note that there is no GSG linker between the first gene and the 2A peptides in these constructs. (B-D) Normalized fluorescence of TagRFP-T and mTurquoise2 in the empty vector, monocistronic control, and experimental bi-cistronic constructs in yeast strains BY4741, CEN.PK2-1C, W303, and CEN.PK2. Data represent the average ± SD of three independent biological replicates.

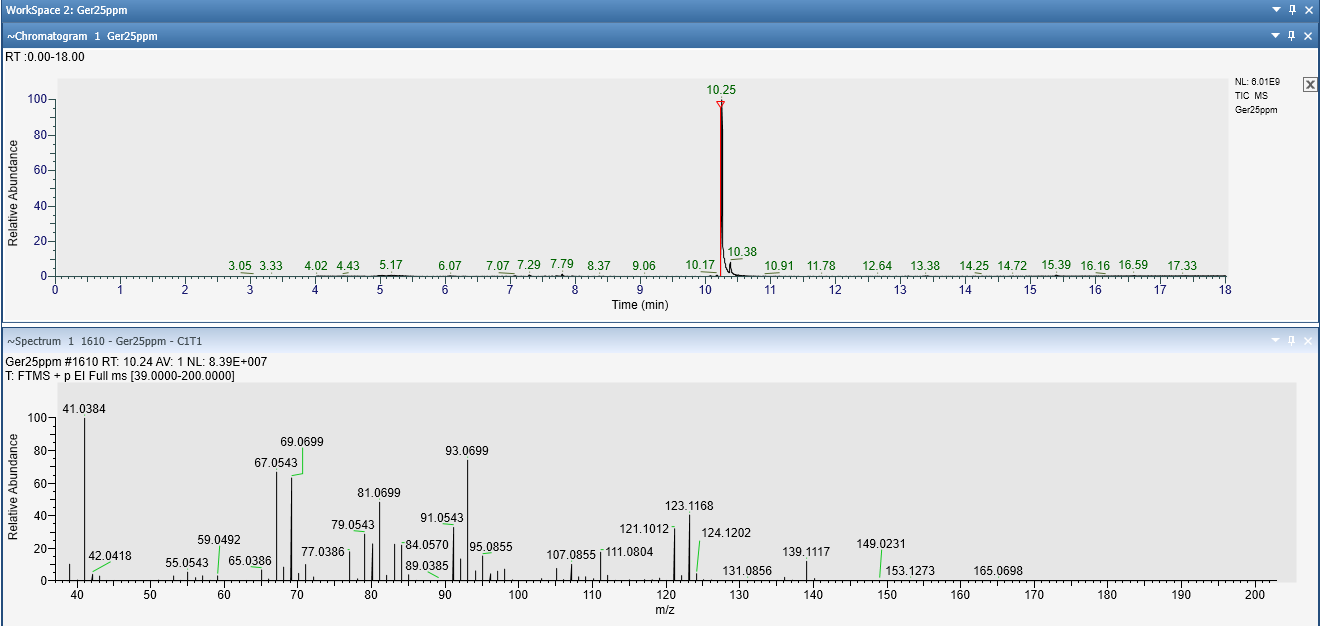

**A**.

**B**

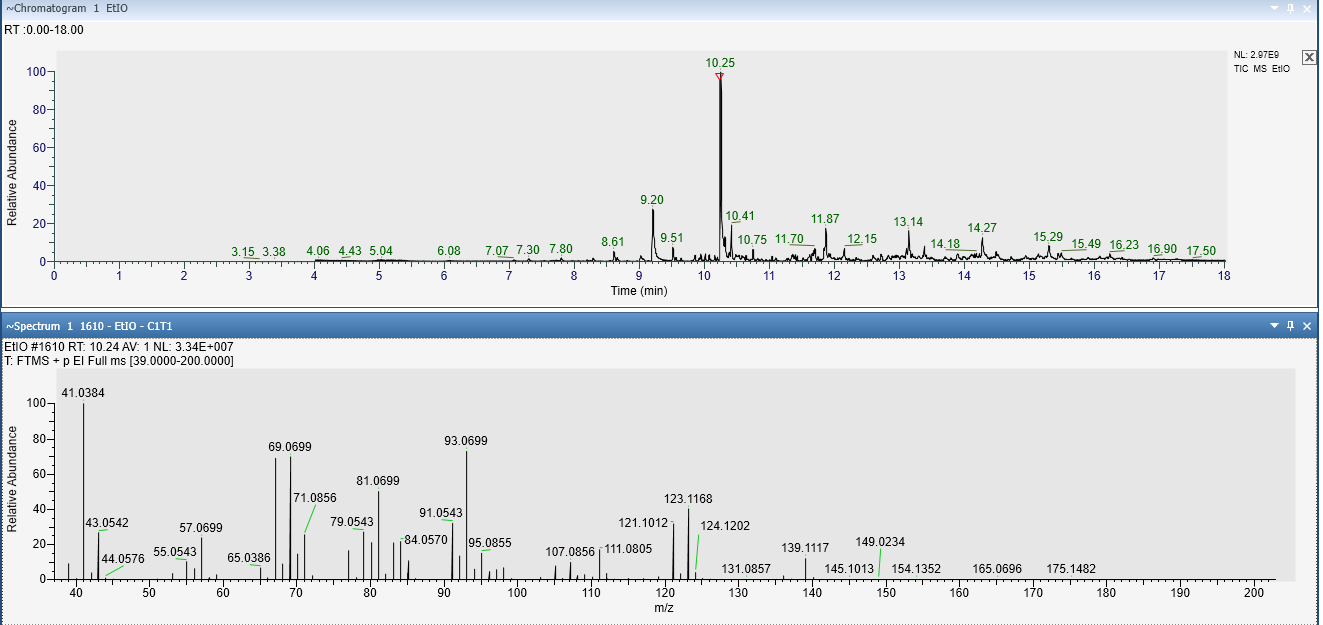

**Figure S9:** GC-MS for geraniol identification. Chromatograms (TIC) and MS spectra of (A) geraniol (25 mg/l) authentic standard, (B) geraniol produced from yeast bearing the 2A_2 construct.

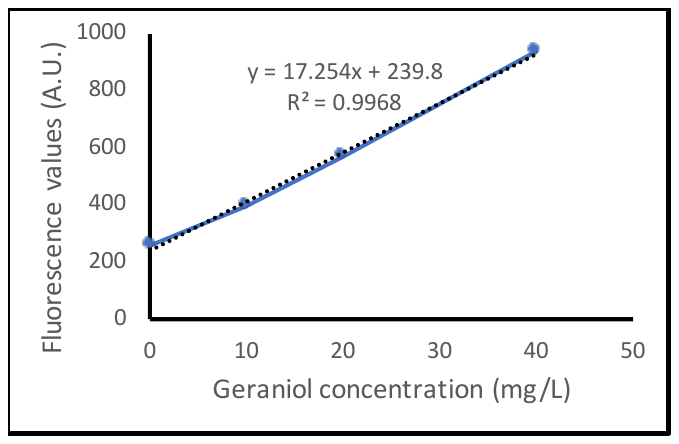

**Figure S10**: Standard curve prepared for the geraniol dehydrogenase assay*^5^* using pure geraniol.

**Table S4:** List of plasmids used in this work. All plasmid names starting in pYTK001 have DNA parts cloned into the entry vector pYTK001. All CDSs have a 5’ overhang with the promoter and a 3’ overhang with the terminator unless otherwise specified. All TUs have the p*GAL1* promoter and the t*CYC1* terminator.

| **Bi-cistronic constructs:** | | |
| --- | --- | --- |
| *Characterizing the four 2A peptides:* | | |
| **Plasmid name** | **Relevant characteristics** | **Source** |
| pYTK001 | Entry vector with GFP dropout to clone DNA parts | Lee et al (2015)*^1^* |
| pFA6a-*link-yoTagRFP-T-Kan* | TagRFP-T was amplified from this plasmid | Lee et al (2013)*^6^* |
| pYTK001_*TagRFP-T-6X His (c)* | TagRFP-T with 6X His tag | This study |
| pYTK032 | pYTK001_*mTurquoise2* from the yeast MolClo kit | Lee et al (2015)*^1^* |
| pYTK001_*mTurquoise2-6X His (c)* | mTurquoise2 with 6X His tag |  |
| pYTK001_*E2Ab* | GSG-E2A 2A cloned into pYTK001 | This study |
| pYTK001_*O2Ab* | GSG-O2A 2A cloned into pYTK001 | This study |
| pYTK001_*P2Ab* | GSG-P2A 2A cloned into pYTK001 | This study |
| pYTK001_*T2Ab* | GSG-T2A 2A cloned into pYTK001 | This study |
| pYTK001_*tCYC1* | tCYC1 terminator cloned in pYTK001 | This study |
| pTU1312 (inter)_*GFP dropout* | Intermediate TU vector with GFP dropout, CEN origin, *HIS3* selection marker | This study |
| pTU1312_*mTurquoise2-6X His (c)* | TU to express mTurquoise2 as control | This study |
| pTU1312_*TagRFP-T-6X His (c)* | TU to express TagRFP-T as control | This study |
| pTU1312_*mTurquoise2.E2A. TagRFP-T-6X His* | Bi-cistronic construct with mTurquoise2 and TagRFP-T separated by E2A | This study |
| pTU1312_mTurquoise2.O2A. TagRFP-T-6X His | Bi-cistronic construct with mTurquoise2 and TagRFP-T separated by O2A | This study |
| pTU1312_*mTurquoise2.P2A. TagRFP-T-6X His* | Bi-cistronic construct with mTurquoise2 and TagRFP-T separated by P2A | This study |
| pTU1312_*mTurquoise2.T2A. TagRFP-T-6X His* | Bi-cistronic construct with mTurquoise2 and TagRFP-T separated by T2A | This study |
| pTU1312_*TagRFP-T.E2A. mTurquoise2-6X His* | Bi-cistronic construct with TagRFP-T and mTurquoise2 separated by E2A | This study |
| pTU1312_*TagRFP-T. O2A. mTurquoise2-6X His* | Bi-cistronic construct with TagRFP-T and mTurquoise2 separated by O2A | This study |
| pTU1312_*TagRFP-T.P2A. mTurquoise2-6X His* | Bi-cistronic construct with TagRFP-T and mTurquoise2 separated by P2A | This study |
| pTU1312_*TagRFP-T.T2A. mTurquoise2-6X His* | Bi-cistronic construct with TagRFP-T and mTurquoise2 separated by T2A | This study |
| *Testing different linker peptides:* | | |
| pYTK001_(*GSG)4-E2A* | (GSG)4-E2A 2A cloned into pYTK001 | This study |
| pYTK001_*Rigid-E2A* | Rigid linker, A(EAAAK)2 fused to E2A | This study |
| pYTK001_*Flexible-E2A* | Flexible linker, GSAGSAAGSGEF fused to E2A | This study |
| pTU1312_*mTurquoise2.(GSG)4-E2A. TagRFP-T-6X His* | Bi-cistronic construct with mTurquoise2 and TagRFP-T separated by (GSG)4-E2A | This study |
| pTU1312_*mTurquoise2.Rigid-E2A. TagRFP-T-6X His* | Bi-cistronic construct with mTurquoise2 and TagRFP-T separated by Rigid-E2A | This study |
| pTU1312_*mTurquoise2.Rigid-E2A. TagRFP-T-6X His* | Bi-cistronic construct with mTurquoise2 and TagRFP-T separated by Flexible-E2A | This study |
| *Testing the effect of E2A fused to fluorescence proteins:* | | |
| pYTK001_*mTurquoise2-E2A* | mTurquoise2 and E2A fusion | This study |
| pYTK001_*TagRFP-T-E2A* | TagRFP-T and E2A fusion | This study |
| pYTK001_*mKate2-E2A* | Venus and E2A fusion | This study |
| pTU1312_*mTurquoise2-E2A* | TU having E2A fused to mTurquoise2 | This study |
| pTU1312_ *TagRFP-T-E2A* | TU having E2A fused to TagRFP-T | This study |
| pTU1312_ *mKate2-E2A* | TU having E2A fused to mKate2 | This study |
| *Testing the effect of E2A fused to lacZ:* | | |
| pYTK001_*lacZ* | lacZ gene from *E. coli* MG1655 | This study |
| pYTK001_*lacZ- E2A* | lacZ gene fused to E2A | This study |
| pTU1312_*lacZ* | TU for expression of lacZ | This study |
| pTU1312_*lacZ- E2A* | TU for expression of lacZ-E2A | This study |
| **Experiments with the tri-cistronic constructs** | | |
| pYTK033 | pYTK001_*Venus* from the yeast MolClo kit | Lee et al (2015)*^1^* |
| pYTK001_*Venus-HA* | Venus with HA tag |  |
| pTU1312_*Venus-HA (c)* | TU to express Venus only as control | This study |
| pYTK001_*TagRFP-T_2A* | TagRFP-T with 3’ overhang for the 1^st^ 2A | This study |
| pYTK001_*2A_mTurquoise2* | mTurquoise2 with 5’ overhang for the 1^st^ 2A and 3’ overhang for the 2^nd^ 2A | This study |
| pYTK001_*2A_Venus-HA* | Venus with 5’ overhang for the 2^nd^ | This study |
| pYTK001_*E2At(1^st^)* | GSG-E2A having 5’ overhang with TagRFP-T and 3’ overhang with mTurquoise2 | This study |
| pYTK001_*O2At(1^st^)* | GSG-O2A having 5’ overhang with TagRFP-T and 3’ overhang with mTurquoise2 | This study |
| pYTK001_*P2At(1^st^)* | GSG-P2A having 5’ overhang with TagRFP-T and 3’ overhang with mTurquoise2 | This study |
| pYTK001_*E2At(2^nd^)* | GSG-E2A having 5’ overhang with mTurquoise2 and 3’ overhang with Venus | This study |
| pYTK001_*O2At(2^nd^)* | GSG-O2A having 5’ overhang with mTurquoise2 and 3’ overhang with Venus | This study |
| pYTK001_*P2At(2^nd^)* | GSG-P2A having 5’ overhang with mTurquoise2 and 3’ overhang with Venus | This study |
| pTU1312(inter)_*pGAL1.GFP dropout.tCYC1* | Intermediate TU level vector with pGAL1 promoter, GFP dropout and tCYC1 terminator | This study |
| pTU1312_*TagRFP-T.E2A.mTurquoise2.O2A.Venus-HA* | Tri-cistronic construct with TagRFP-T, mTurquoise2 and Venus separated by E2A and O2A. | This study |
| pTU1312_*TagRFP-T. O2A.mTurquoise2.E2A.Venus-HA* | Tri-cistronic construct with the same fluorescence proteins separated by O2A and E2A | This study |
| pTU1312_*TagRFP-T.E2A.mTurquoise2.P2A.Venus-HA* | Tri-cistronic construct with the same fluorescence proteins separated by E2A and P2A | This study |
| pTU1312_*TagRFP-T.P2A. mTurquoise2.E2A.Venus-HA* | Tri-cistronic construct with the same fluorescence proteins separated by P2A and E2A | This study |
| pTU1312_*TagRFP-T.P2A. mTurquoise2.O2A.Venus-HA* | Tri-cistronic construct with the same fluorescence proteins separated by P2A and O2A | This study |
| pTU1312_*TagRFP-T.O2A. mTurquoise2.P2A.Venus-HA* | Tri-cistronic construct with the same fluorescence proteins separated by O2A and P2A | This study |
| **Experiments with the fluorescence quad-cistronic construct** | | |
| pFA6a-*link-yomKate2-SpHis5* | mKate2 was amplified from this plasmid | Lee et al (2013)*^6^* |
| pYTK001_*mKate2 (c)* | mKate2 cloned in pYTK001 | This study |
| pTU1312_*mKate2 (c)* | TU to express mKate2 as control | This study |
| pYTK001_*mKate2_2A* | mKate2 with 3’ overhang for the 1^st^ 2A peptide | This study |
| pYTK001_*Venus-6X His (c)* | Venus with a 6X His tag |  |
| pYTK001_*2A_TagRFP-T* | TagRFP-T with 5’ overhang for the 1^st^ 2A and 3’ overhang for the 2^nd^ 2A | This study |
| pYTK001_*2A_mTurquoise2* | mTurquoise2 with 5’ overhang for the 2^nd^ 2A and 3’ overhang for the 3^rd^ 2A | This study |
| pYTK001_*2A_Venus-6X His* | Venus-6X His with 5’ overhang for the 3^rd^ 2A | This study |
| pYTK001_*E2Aq(1^st^)* | GSG-ERBV with 5’ overhang for mKate2 and 3’ overhang for TagRFP-T | This study |
| pYTK001_*P2Aq(2^nd^)* | GSG-P2A with 5’ overhang for TagRFP-T and 3’ overhang for mTurquoise2 | This study |
| pYTK001_*O2Aq (3^rd^)* | Same as pYTK001_*O2At* (2^nd^) of the tri-cistronic constructs | This study |
| pTU1312_*mKate2.E2A.TagRFP-T.P2A.mTUrquoise2.O2A.Venus-6X His* | Quad-cistronic construct for the co-expression of mKate2, TagRFP-T, mTurquoise2 and Venus | This study |
| **Expression of the MVA pathway** | | |
| *Control Multigene construct:* | | |
| pYTK001_*tHMG1* | *tHMG1* from the yeast genome | This study |
| pYTK001_*ERG20^ww^* | *ERG20^ww^* , a mutant version of *ERG20* | This study |
| pYTK001_*IDI1* | *IDI1* from the yeast genome | This study |
| pYTK001_*tObGES* | *Truncated Ob*GES from *Ocimum basilicum* | This study |
| pTU1312_*tHMG1* | TU to express *tHMG1* | This study |
| pTU1312_*ERG20^ww^* | TU to express *ERG20^ww^* | This study |
| pTU1312_*IDI1* | TU to express *IDI1* | This study |
| pTU1312_*tOb*GES | TU to express t*Ob*GES | This study |
| pMGR13_*tHMG1. ERG20^ww^. IDI1.* *tOb*GES | Multigene plasmid to express *tHMG1, ERG20^ww^, IDI1* and *tOb*GES separately. | This study |
| *Experimental 2A constructs:* | | |
| pYTK001_*tHMG1* (1^st^) | *tHMG1* with 3’ overhang for E2A | This study |
| pYTK001_*tHMG1* (2^nd^) | *tHMG1* with 5’ overhang for E2A and 3’ overhang for P2A | This study |
| pYTK001_*tHMG1* (3^rd^) | *tHMG1* with 5’ overhang for P2A and 3’ overhang for O2A | This study |
| pYTK001_*tHMG1* (4^th^) | *tHMG1* with 5’ overhang for O2A | This study |
| pYTK001_*ERG20^ww^* (1^st^) | *ERG20^ww^* with 3’ overhang for E2A | This study |
| pYTK001_*ERG20^ww^* (2^nd^) | *ERG20^ww^* with 5’ overhang for E2A and 3’ overhang for P2A | This study |
| pYTK001_*ERG20^ww^* (3^rd^) | *ERG20^ww^* with 5’ overhang for P2A and 3’ overhang for O2A | This study |
| pYTK001_*ERG20^ww^* (4^th^) | *ERG20^ww^* with 5’ overhang for O2A | This study |
| pYTK001_*tOb*GES (1^st^) | t*Ob*GES with 3’ overhang for E2A | This study |
| pYTK001_*tOb*GES (4^th^) | t*Ob*GES with 5’ overhang for O2A | This study |
| pYTK001_*IDI1* (2^nd^) | *IDI1* with 5’ overhang for E2A and 3’ overhang for P2A | This study |
| pYTK001_*IDI1* (3^rd^) | *IDI1* with 5’ overhang for P2A and 3’ overhang for O2A | This study |
| pYTK001_*IDI1* (4^th^) | *tHMG1* with 5’ overhang for O2A | This study |
| pTU1312_*tHMG1.E2A. ERG20^ww^.P2A.IDI1.O2A.tObGES* | Quad-cistronic construct with *tHMG1,* *ERG20^ww^, IDI1* and *tOb*GES in order. | This study |
| pTU1312_*ERG20^ww^. E2A. tHMG1. P2A.IDI1.O2A.tObGES* | Quad-cistronic construct with *ERG20^ww^*, *tHMG1, IDI1* and *tOb*GES in order. | This study |
| pTU1312_*tObGES.E2A.IDI1.P2A. tHMG1.O2A. ERG20^ww^* | Quad-cistronic construct with *tOb*GES, *IDI1*, *tHMG1* and *ERG20^ww^* in order. | This study |
| pTU1312_*tObGES.E2A.IDI1.P2A. ERG20^ww^. O2A. tHMG1* | Quad-cistronic construct with *tOb*GES, *IDI1*, *ERG20^ww^* and *tHMG1* in order. | This study |

**Table S5:** List of primers used in this work. Key: BsaI overhang BsmbI overhang

| **Primers** | **Sequence (5’-3’)** | **Description** |
| --- | --- | --- |
| **Bi-cistronic constructs** | | |
| TagRFP-T F | tttcgtctcgtcggtctcatatggtatctaaaggt  gaagagttg | Forward primer to amplify TagRFP-T with 5’ overhang for the promoter |
| 2A_TagRFP-T F | tttcgtctcgtcggtctcgcctatggtatctaa  aggtgaagagttg | Forward primer to amplify TagRFP-T with 5’ overhang for the 2A peptide. |
| TagRFP-T-6X His R | tttcgtctcgggtctcaggatttagtgatgatgatg  atgatgcttatacaattcatccataccattcag | Reverse primer to amplify TagRFP-T-6X His with 3’ overhang for the terminator |
| TagRFP-T_2A R | tttcgtctcgggtcggtctcagaacccttatacaat  tcatccataccattcag | Reverse primer to amplify TagRFP-T with 3’ overhang for the 2A peptide |
| mTurquoise2 F | tttcgtctcgtcggtctcatatggtttctaaag  gtgaagaattattc | Forward primer to amplify mTurquoise2 with 5’ overhang for the promoter |
| 2A_mTurquoise2 F | tttcgtctcgtcggtctcacctatggtttct  aaaggtgaagaattattc | Forward primer to amplify TagRFP-T with 5’ overhang for the 2A peptide |
| mTurquoise2-6X His R | tttcgtctcgggtctcaggatttagtgatgatg  atgatgatgtttgtacaattcatccatacccaag | Reverse primer to amplify mTurquoise2-6X His with 3’ overhang for the terminator |
| mTurquoise2_2A R | tttcgtctctggtctcagaactttgtacaatt  catccatacccaag | Reverse primer to amplify mTurquoise2 with 3’ overhang for the 2A peptide |
| *Primers to amplify 2A peptides, applicable for bi-cistronic and the 1^st^ 2A peptide of tri-cistronic constructs* | | |
| GSG-E2A F | tttcgtctcgtcggtctcagttctggtggcgctac | Forward primer to amplify GSG-E2A with 5’ overhang for the 1^st^ gene |
| GSG-E2A R | tttcgtctcgggtctcatagggccgggattcaattc | Reverse primer to amplify GSG-E2A with 3’ overhang for the 2^nd^ gene |
| GSG-O2A F | tttcgtctcgtcggtctcagttctggtaagtccaattacg | Forward primer to amplify GSG-O2A with 5’ overhang for the 1^st^ gene |
| GSG-O2A R | tttcgtctcgggtctcataggtcctgggttggattc | Reverse primer to amplify GSG-O2A with 3’ overhang for the 2^nd^ gene |
| GSG-P2A F | tttcgtctcgtcggtctcagttctggtgcaacaaatttc | Forward primer to amplify GSG-P2A with 5’ overhang for the 1^st^ gene |
| GSG-P2A R | tttcgtctcgggtctcataggtccagggttctcctccacatc | Reverse primer to amplify GSG-P2A with 3’ overhang for the 2^nd^ gene |
| GSG-T2A F | tttcgtctcgtcggtctcagttctggtgaaggtaggggctc | Forward primer to amplify GSG-T2A with 5’ overhang for the 1^st^ gene |
| GSG-T2A R | tttcgtctcgggtctcataggtcctggattctcttcaac | Reverse primer to amplify GSG-T2A with 3’ overhang for the 2^nd^ gene |
| tCYC1 F | tttcgtctcgtcggtctcaatccgctctaaccgaaaagg | Forward primer to amplify tCYC1 with 5’ overhang for the last gene |
| tCYC1 R | tttcgtctcgggtctcacagccttcgagcgtcccaaaac | Reverse primer to amplify tCYC1 with 3’ overhang for the right connector |
| (GSG)4-E2A F | tttcgtctcgtcggtctcgcaaagggtctggtggctcag  gagggagtggtggttctggtggcgctaccaacttctc | Forward primer having the (GSG)X4 sequence to amplify E2A with 5’ overhang for mTurquoise2 |
| Rigid-E2A F | tttcgtctcgtcggtctcgcaaagcggaggcagcggcg  aaagaagccgccgccaaagcgggcgctaccaatttctc | Forward primer having the A(EAAAK)X2 sequence to amplify E2A with 5’ overhang for mTurquoise2 |
| Flexible E2A F | tttcgtctcgtcggtctcacaaagggagcgcgggcagt  gcagccgggtcaggcgagttcggcgctaccaacttctc | Forward primer having the GSAGSAAGSGEF sequence to amplify E2A with 5’ overhang for mTurquoise2 |
| mTurquoise2_E2A R | tttcgtctcgatcctttgtacaattcatccatacccaag | Reverse primer to amplify mTurquoise2 with 3’ overhang for E2A |
| TagRFP-T_E2A R | tttcgtctcgatcccttatacaattcatccataccattcag | Reverse primer to amplify TagRFP-T with 3’ overhang for E2A |
| mKate2_E2A R | tttcgtctcgatcctctgtgtcccaacttagatgg | Reverse primer to amplify mKate2 with 3’ overhang for E2A |
| Fl. Pr.+ E2A F | tttcgtctcgggatccggcgctacc | Forward primer to amplify GSG-E2A with 5’ overhang for fluorescence genes |
| ERBV+ stop R | tttcgtctctggtctctggattcagccgggattcaattcgac | Reverse primer with stop codon to amplify GSG-E2A with 3’ overhang for terminator. |
| lacZ Gb F | caaggagaaaaaaccccggatccatgaccatgattacgg  attcac | Forward primer to amplify lacZ with overhang for Gibson cloning |
| lacZ Gb R | agccgcggtaccaagcttactcgagtcatttttgacaccag  accaactg | Reverse primer to amplify lacZ with overhang for Gibson cloning |
| lacZ_GSG-E2A Gb R | agcggtaccaagcttactcgagttagccgggattcaattcg  acgtcacccgccagtttcaatagtgagaagttggtagcgcc  accagaacctttttgacaccagaccaac | Reverse primer to amplify lacZ with GSG-E2A and overhang for Gibson cloning |
| **Tri-cistronic constructs** | | |
| *TagRFP-T F and TagRFP-T_2A are the same as in bi-cistronic constructs* | | |
| 2A_mTurquoise2 F | tttcgtctcgtcggtctcgcctatggtttctaaaggtgaaga  attattc | Forward primer to amplify mTurquoise2 with 5’ overhang for the 1^st^ 2A peptide |
| 2A_mTurquoise2 R | Tttcgtctcgggtctctcggaacctttgtacaattcatccata  cccaag | Reverse primer to amplify mTurquoise2 with 3’ overhang for the 2^nd^ 2A peptide |
| Venus (c) F | tttcgtctcgtcggtctcgtatgtctaaaggtgaaga  attattcactg | Forward primer to amplify Venus with 5’ overhang for the promoter |
| 2A_Venus F | Tttcgtctcgtcggtctcgcaggacctatgtctaaaggtgaagaattattcactg | Forward primer to amplify Venus with 5’ overhang for the 2^nd^ 2A peptide |
| Venus-HA R | tttcgtctcggtctctcgggattcaagcgtaatctggaa  catcgtatgggtaggatcctttgtacaattcatccata | Reverse primer to amplify Venus with HA tag and overhang with terminator |
| E2A 2^nd^ F | tttcgtctcgtcggtctcgtccggtggcgctacc | Forward primer to amplify GSG-E2A with 5’ overhang with the 2^nd^ gene |
| E2A 2^nd^ R | tttcgtctcgggtcggtctcccctggattcaattcgacgtcac | Reverse primer to amplify GSG-E2A with 3’ overhang with the 3^rd^ gene |
| O2A 2^nd^ F | tttcgtctcgtcggggtctcttccggtaagtccaattacgac | Forward primer to amplify GSG-O2A with 5’ overhang with the 2^nd^ gene |
| O2A 2^nd^ R | tttcgtctcgggtctcccctgggttggattcaacatctc | Reverse primer to amplify GSG-O2A with 3’ overhang with the 3^rd^ gene |
| P2A 2^nd^ F | tttcgtctcgtcggtctcgtccggtgcaacaaatttctcattg | Forward primer to amplify GSG-P2A with 5’ overhang with the 2^nd^ gene |
| P2A 2^nd^ R | tttcgtctcgggtctcccctgggttctcctccacatc | Reverse primer to amplify GSG-P2A with 3’ overhang with the 3^rd^ gene |
| **Fluorescence Quad-cistronic construct** | | |
| mKate2 F | tttcgtctcgtcggtctcgtatggtttctgaactcatcaagg | Forward primer to amplify mKate2 with 5’ overhang with the promoter |
| mKate2 (c) R | tttcgtctcgggtctccggatttatctgtgtcccaacttaga  tgg | Reverse primer to amplify mKate2 with 3’ overhang with the terminator |
| mKate2_2A R | tttcgtctcgggtctccgctccctctgtgtcccaacttagatgg | Reverse primer to amplify mKate2 with 3’ overhang with the 1^st^ 2A peptide |
| 2A_TagRFP-T F | tttcgtctcgtcggtctcgcctatggtatctaaaggtgaag  agttg | Forward primer to amplify TagRFP-T with 5’ overhang with the 1^st^ 2A peptide |
| 2A_TagRFP-T R | tttcgtctcgggtctcggaacccttatacaattcatccatac  cattcag | Reverse primer to amplify TagRFP-T with 3’ overhang with the 2^nd^ 2A peptide |
| 2A_mTurquoise2 F | tttcgtctcgtcggtctcgaggacctatggtttctaaaggtg  aagaattattc | Forward primer to amplify mTurquoise2 with 5’ overhang with the 2^nd^ 2A peptide |
| 2A_mTurquoise2 R | tttcgtctcgggtctcccggaacctttgtacaattcatccat  acccaag | Reverse primer to amplify mTurquoise2 with 3’ overhang with the 3^rd^ 2A peptide |
| E2A 1^st^ F | tttcgtctcgtcggtctcggagcggtggagctaccaacttctc | Forward primer to amplify GSG-E2A with 5’ overhang with the 1^st^ gene |
| E2A 1^st^ R | tttcgtctcgggtctcctagggccgggattcaattc | Reverse primer to amplify GSG-E2A with 3’ overhang with the 2^nd^ gene |
| P2A 2^nd^ F | tttcgtctcgtcggtctcggttctggtgcaacaaatttc | Forward primer to amplify GSG-P2A with 5’ overhang with the 2^nd^ gene |
| P2A 2^nd^ R | tttcgtctcgggtctcctcctgggttctcctccacatctc | Reverse primer to amplify GSG-P2A with 3’ overhang with the 3^rd^ gene |
| O2A 3^rd^ F and R and 2A_Venus F for quad-cistronic construct are same as O2A 2^nd^ F and R and 2A_Venus F of the tri-cistronic constructs. | | |
| **MVA pathway (Quad-cistronic constructs)** | | |
| tHMG1 F | tttcgtctcgtcggtctcgtatgccagttttaaccaataaa  acag | Forward primer to amplify tHMG1 with 5’ overhang for the promoter |
| tHMG1 R | tttcgtctctggtctccggatttaggatttaatgcaggtga  cgg | Reverse primer to amplify tHMG1 with 3’ overhang for the terminator |
| tHMG1 1^st^ R | tttcgtctcgggtctctgctcccggatttaatgcaggtg  acgg | Reverse primer to amplify tHMG1 with 3’ overhang for the 1^st^ 2A peptide |
| tHMG1 2^nd^ F | tttcgtctcgtcggtctcgcctatgccagttttaaccaat  aaaacag | Forward primer to amplify tHMG1 with 5’ overhang for the 1^st^ 2A peptide |
| tHMG1 2^nd^ R | tttcgtctcaggtctcggaaccggatttaatgcaggtg  acgg | Reverse primer to amplify tHMG1 with 3’ overhang for the 2^nd^ 2A peptide |
| tHMG1 3^rd^ F | tttcgtctcgtcggtctcaaggacctatgccagttttaa  ccaataaaacag | Forward primer to amplify tHMG1 with 5’ overhang for the 2^nd^ 2A peptide |
| tHMG1 3^rd^ R | tttcgtctcaggtcggtctctcggaaccggatttaatgc  aggtgacgg | Reverse primer to amplify tHMG1 with 3’ overhang for the 3^rd^ 2A peptide |
| tHMG1 4^th^ F | tttcgtctcgtcggtctcgcaggacctatgccagttttaa  ccaataaaaca | Forward primer to amplify tHMG1 with 5’ overhang for the 3^rd^ 2A peptide |
| ERG20^ww^ F | tttcgtctcgtcggggtctcgtatggcttcagaaaaaga  aattagg | Forward primer to amplify ERG20^ww^ with 5’ overhang for the promoter |
| ERG20^ww^ R | tttcgtctctggtctccggatctatttgcttctcttgtaaa  ctttgttc | Reverse primer to amplify ERG20^ww^ with 3’ overhang for the terminator |
| ERG20^ww^ 1^st^ R | tttcgtctctggtctcggctccctttgcttctcttgtaaac  tttgttc | Reverse primer to amplify ERG20^ww^ with 3’ overhang for the 1^st^ 2A peptide |
| ERG20^ww^ 2^nd^ F | tttcgtctcgtcggtctcgcctatggcttcagaaaaaga  aattaggag | Forward primer to amplify ERG20^ww^ with 5’ overhang for the 1^st^ 2A peptide |
| ERG20^ww^ 2^nd^ R | tttcgtctctggtctctgaacctttgcttctcttgtaaact  ttgttc | Reverse primer to amplify ERG20^ww^ with  3’ overhang for the 2^nd^ 2A peptide |
| ERG20^ww^ 3^rd^ F | tttcgtctcgtcggtctcgaggacctatggcttcagaaaa  agaaattaggag | Forward primer to amplify ERG20^ww^ with 5’ overhang for the 2^nd^ 2A peptide |
| ERG20^ww^ 3^rd^ R | tttcgtctctggtctctcggaacctttgcttctcttgtaaac  tttgttc | Reverse primer to amplify ERG20^ww^ with  3’ overhang for the 3^rd^ 2A peptide |
| ERG20^ww^ 4^th^ F | tttcgtctcgtcggtctcgcaggacctatggcttcagaaa  aagaaattaggag | Forward primer to amplify ERG20^ww^ with 5’ overhang for the 3^rd^ 2A peptide |
| IDI1 F | tttcgtctcgtcggtctcgtatgactgccgacaacaatag | Forward primer to amplify IDI1 with 5’ overhang for the promoter |
| IDI1 R | tttcgtctctggtctccggatttatagcattctatgaatttg  cctgtc | Reverse primer to amplify IDI1 with 3’ overhang for the terminator |
| IDI1 2^nd^ F | tttcgtctcgtcggtctcgcctatgactgccgacaacaa  tag | Forward primer to amplify IDI1 with 5’ overhang for the 1^st^ 2A peptide |
| IDI1 2^nd^ R | tttcgtctcaggtctctgaacctagcattctatgaatttgc  ctgtc | Reverse primer to amplify IDI1 with 3’ overhang for the 2^nd^ 2A peptide |
| IDI1 3^rd^ F | tttcgtctcttcggtctcgaggacctatgactgccgacaac  aatag | Forward primer to amplify IDI1 with 5’ overhang for the 2^nd^ 2A peptide |
| IDI1 3^rd^ R | tttcgtctctggtctcccggaacctagcattctatgaatttg  cctgtc | Reverse primer to amplify IDI1 with 3’ overhang for the 3^rd^ 2A peptide |
| tObGES F | tttcgtctcgtcggtctcgtatggaagagagttcatcaaagc | Forward primer to amplify tObGES with 5’ overhang for the promoter |
| tObGES R | tttcgtctcgggtctctggatttattgtgtaaaaaacaggg  catc | Reverse primer to amplify tObGES with 3’ overhang for the terminator |
| tObGES 1^st^ R | Tttcgtctctggtctctgctcccttgtgtaaaaaacagggc  atcg | Forward primer to amplify tObGES with 5’ overhang for the 1^st^ 2A peptide |
| tObGES 4^th^ F | Tttcgtctcgtcggtctcgcaggacctatggaagagagttca  tcaaagc | Reverse primer to amplify tObGES with 3’ overhang for the 3^rd^ 2A peptide |
